# A BRRF1-CCR4-NOT axis underlies conserved transcriptome-wide loss of splicing fidelity during gammaherpesvirus reactivation

**DOI:** 10.64898/2026.05.20.726682

**Authors:** Trang T Nguyen, Anandita Ghosh, Nadeeshika Wickramarachchige-Dona, Tina M O’Grady, Claire Roberts, Eman Ishaq, Jordan Bass, Meggie Lam, Melody Baddoo, Nathan A Ungerleider, Nick Van Otterloo, Jia Wang, Qian Zhang, Hong Liu, Yan Dong, Rolf Renne, Truong D Nguyen, Erik K Flemington

## Abstract

Gammaherpesvirus reactivation drives a collapse of host mRNA splicing fidelity that extends across the transcriptome, with exon skipping affecting up to ∼57% of expressed genes, exceeding the effects of depletion of any of 186 splicing factors. Combining five Epstein-Barr virus (EBV) and Kaposi’s sarcoma-associated herpesvirus (KSHV) reactivation systems across B cell and epithelial models with deep poly(A)+ RNA sequencing of purified lytic cells, we find that most induced isoforms are predicted to undergo nonsense-mediated decay or to lose conserved protein domains, broadly compromising cell cycle, innate immune and RNA-processing pathways. The phenotype arises independently of viral DNA replication, indicating early host remodeling. A screen of EBV early genes identifies BRRF1 as a key driver: through a CIY(Y/E) motif conserved in KSHV ORF49, BRRF1 engages the nuclear CCR4-NOT complex through its CNOT9 and CNOT1 subunits, hijacking this canonically cytoplasmic deadenylation hub for nuclear disruption of host splicing.

## Introduction

Productive gammaherpesvirus replication requires rapid host takeover. To complete the lytic program, the virus must reconfigure host gene-expression pathways to favor viral output while suppressing antiviral responses. Prior work has shown that Epstein-Barr virus (EBV) lytic reactivation remodels the host through mRNA degradation, suppression and redirection of host transcription, and large-scale changes in chromatin organization and nuclear architecture (2–4). A small number of studies have also linked individual lytic factors, including BMLF1 (5,6), to altered splicing of selected host transcripts, and recent RNA sequencing data from reactivating EBV-infected cells suggest that host splicing changes may be broader than previously appreciated (2). However, it remains unclear whether these observations reflect isolated locus-specific effects or a more global failure of host splicing fidelity during reactivation. Such a mechanism would provide an especially efficient mode of host takeover, because widespread disruption of pre-mRNA splicing could simultaneously cripple inducible host defense pathways and erode productive cellular gene output across much of the transcriptome.

EBV provides an important system in which to address this question. EBV is a ubiquitous human oncogenic gammaherpesvirus associated with approximately 250,000 cases of lymphoma and carcinoma annually, and it is also the major cause of infectious mononucleosis and a major environmental risk factor for multiple sclerosis (7–12). Following primary infection, EBV establishes lifelong latency primarily in memory B cells, where a restricted viral gene-expression program preserves the episome while minimizing immune recognition (13–15). In response to appropriate stimuli, EBV can transition from latency into the lytic phase, during which viral DNA is amplified, and progeny virions are produced. Although latency has long been viewed as the dominant viral state in EBV-associated malignancy (16,17), a growing body of evidence indicates that lytic reactivation also contributes to pathogenesis by promoting viral spread and by actively reshaping host-cell programs relevant to disease (18–20).

Recent work has made clear that herpesvirus-mediated host remodeling is not limited to changes in RNA abundance alone. One well-established strategy is host shutoff, in which cellular mRNAs are degraded to reduce competition for translational capacity and favor viral protein synthesis (21,22). In EBV, the viral exonuclease BGLF5 contributes to this process, although recent studies indicate that BGLF5 alone cannot account for the full extent of host transcriptome restructuring observed during reactivation (2,23,24). Additional layers of host takeover have also emerged, including widespread suppression of host transcription through altered chromatin and nuclear architecture and large-scale remodeling of host transcription initiation landscapes (3,4,25). Together, these findings point to a multilayered host-remodeling program and raise the possibility that disruption of host RNA processing constitutes a major arm of the lytic host-takeover strategy.

Among the potential layers of host takeover, disruption of host pre-mRNA splicing may be especially consequential. Splicing is required for productive expression of a large fraction of the human transcriptome, and a broad loss of splicing fidelity would be expected to impair both constitutive cellular functions and newly induced host-response programs. This may be particularly important for antiviral, inflammatory, stress-response, and cell cycle pathways that must be rapidly mobilized during reactivation, because these responses depend on the timely production of correctly processed transcripts. Such a defect could create a favorable asymmetry during lytic replication, as host gene expression is highly dependent on accurate splicing, whereas most EBV lytic genes are mono-exonic and comparatively insulated from splicing defects. Yet despite the central importance of splicing to mammalian gene expression, it remains unresolved whether gammaherpesvirus reactivation induces selected splicing alterations at individual loci or instead drives a broader, more penetrant disruption of host splicing fidelity.

Here, using five complementary reactivation systems spanning EBV and Kaposi’s sarcoma-associated herpesvirus (KSHV), B cell and epithelial contexts, and high-depth RNA sequencing of purified lytic populations, we identify transcriptome-wide loss of host splicing fidelity as a conserved feature of gammaherpesvirus reactivation. This phenotype is dominated by exon skipping and affects more than half of expressed genes, with many resulting isoforms predicted to be nonproductive. We further show that this disruption is established early in the lytic cascade and identify the EBV early protein BRRF1 as a key driver. Mechanistically, BRRF1 engages the CCR4-NOT complex in the nucleus, and depletion of CNOT1 and CNOT9 attenuates exon skipping during reactivation. Together, these findings define a BRRF1-CCR4-NOT axis underlying widespread collapse of host RNA-processing fidelity.

## Materials and Methods

### Generation of stable pCEP4-BMRF1p-GFP lytic reporter cell lines

EBV-positive Burkitt’s lymphoma cell lines Mutu (provided by Samuel H. Speck), Akata (provided by Kenzo Takada), and the EBV-positive stomach cancer cell line SNU719 (provided by the Korean Cell line Bank), were transfected via nucleofection with the episomally replicating lytic reporter containing BMRF1p-driven GFP (pCEP4-BMRF1p-GFP) and a hygromycin-resistant gene. Transfected cells were selected in media supplemented with 250 µg/mL hygromycin (Thermo Scientific, Cat. No. 10687010) for at least 7 days prior to use.

### Cell culture

Mutu, Akata, TREX-BCBL1 (provided by Rolf Renne and Jae Jung), SNU719, and the EBV-negative Burkitt’s lymphoma cell line DG75 (ATCC, Cat. No. CRL-2625) were maintained in RPMI medium (Fisher Scientific, Cat. No. SH30027) supplemented with 10% FBS (Thermo Fisher Scientific, Cat. No. 10437). Akata-BMRF1p-GFP, Mutu-BMRF1p-GFP, and SNU719-BMRF1p-GFP cells were cultured in RPMI + 10% FBS media supplemented with hygromycin B (250 µg/mL). All cells were cultured at 37°C in a 5% CO_2_ incubator.

### Nucleofection

For Zta induction experiments, Mutu cells (3 million) were transfected with 0.3 µg of a CMVp-GFP expression vector plus 3.0 µg of either the SV40p-Zta or control SV40p-Cntl vectors (N=3 per group). BMRF1p-GFP-SNU719 cells (3 million) were transfected with 5 µg of the SV40p-Zta and 5 µg the pLVX-Rta expression vectors or 5 µg of SV40p-Cntl and 5 µg of the pLVX-Cntl control vectors (N=3 per group). DG75 cells (3 million) were transfected with 0.3 µg of the CMVp-GFP expression vector plus either 3 µg of pLVX-expression vectors containing each of the EBV early genes or 3.0 µg of the pLVX-Cntl control vector (N=3 or N=4 per group). Transfections were performed using 100 µL of Amaxa Cell Line Nucleofector Kit R (Lonza, Cat. No. VCA-1001) with the Amaxa Nucleofector II machine (Lonza). Cells, plasmids, and Nucleofector R mixtures were electroporated in a cuvette using pulse code G016. Cells were subsequently transferred to a 6-well plate containing pre-warmed media and incubated for 24 hours before harvesting for FACS sorting.

### BCR crosslinking

Mutu-BMRF1p-GFP and Akata-BMRF1p-GFP cells (3 million) were treated with media containing either no or goat α-Human IgM (Sigma-Aldrich, Cat. No. 10759) (Mutu-BMRF1p-GFP cells) or Affinipure Goat α-Human IgG (Jackson ImmunoResearch, Cat. No. 109-005-003) (Akata-BMRF1p-GFP cells) at a final concentration of 10 µg/mL (N=3 per group). Cells were incubated for 24 hours before being collected for FACS sorting.

### Doxycycline-induced reactivation of KSHV BCBL1 cells

TREx BCBL1-Rta cells (3 million) were resuspended in RPMI supplemented with 15% FBS, 100 µg/mL hygromycin, 50 µg/mL blasticidin (A.G. Scientific, Cat. No B-1247-SOL), and either DMSO for controls (Sigma-Aldrich, Cat. No. D2650-5X5ML) or 1 µg/mL of doxycycline (Sigma-Aldrich, Cat. No. D3072-1ML) (N=3 per group). Cells were then transferred into a 6-well plate and incubated for 24 hours before harvesting.

### Phosphonoacetic acid (PAA) treatment in reactivation experiments

Mutu cells (3 million) were co-transfected with 0.3 µg of the CMVp-GFP expression vector and 3.0 µg of either the empty vector control, SV40p-Cntl or the Zta expression vector, SV40p-Zta by nucleofection, followed by incubation for 24 hours in RPMI + 10% FBS media with either no phosphonoacetic acid or 200 µg/mL phosphonoacetic acid (PAA) (Sigma-Aldrich, Cat. No. 284270-10G) (N=4 per group). Transfected cells were then subjected to FACS sorting for GFP-positive (transfected) cells in PAA-treated and -untreated conditions.

### Fluorescence-activated cell sorting

Cells were harvested and transferred into 15mL tubes and then centrifuged at 1500 rpm for 5 minutes. After aspirating the media, cell pellets were resuspended in DPBS (Gibco, Cat. No. 14190144) and passed through a 35µm mesh filter (Genesee Scientific, Cat. No. 28-154). GFP+ or GFP-cells were then isolated using a BD FACSAria III (BD Biosciences) sorter. Sorted cells were centrifuged again at 1500 rpm for 5 minutes, and the resulting cell pellets were frozen at −80°C prior to RNA extraction.

### RNA extraction and cDNA synthesis

Cell pellets were extracted using TRIzol Reagent (Invitrogen, Cat. No. 15596026) or the RNeasy Mini Kit (Qiagen, Cat. No. 74106) following the manufacturer’s protocol. After TRIzol extractions, RNA pellets were reconstituted in 42 µL of ddH₂O. Five µL of 10× DNase buffer and 3 µL of recombinant DNase I (New England Biotechnology, Cat. No. M0303L) were added to each sample and samples were then incubated at 37°C for 15 minutes. A Monarch RNA Cleanup Kit (New England Biotechnology (Cat. No. T2030L)) was then used to remove DNase I. For RNA preparations using the RNeasy Mini Kit, RNA pellets were reconstituted in 30-50 µL of ddH₂O. Residual DNA was digested using the RNase-Free DNase Set (Qiagen, Cat. No. 79256) according to the manufacturer’s protocol.

After isolation, RNA quality was assessed using the Agilent 2100 Bioanalyzer (Agilent, Serial No. DE54107860) with the Agilent RNA 6000 Nano Kit (Agilent, Cat. No. 5067-1511). RNAs were then subjected to RNA sequencing or used for PCR or qPCR reactions. For PCR reactions, 200 ng to 1 µg of RNA was reverse transcribed using the iScript cDNA Synthesis Kit (Bio-Rad, Cat. No. 1708891) following the manufacturer’s protocol.

### PCR validation of alternative splicing

cDNAs were diluted 1:10 in ddH2O. PCR reactions were carried out using 10 µL of GoTaq Green Mastermix (Promega, cat. no. M7122), 2 µL of diluted cDNA, 0.5 µL of 10µM primer mix, and 7.5 µL of ddH2O (for a final reaction volume of 20 µL). Primer sequences for POLR2A and PLCB3 are provided in **Supplemental Table S3**. PCR was performed using an initial melting step at 95°C for 2 minutes followed by 35-40 PCR cycles (36 cycles for POLR2A and 40 cycles for PLCB3) of 95°C for 30s, 60°C for 30s, and 72°C for 60s. A final extension was then carried out at 72°C for 5 minutes. The PCR products were then run on a 1% agarose gel containing ethidium bromide and imaged using a gel documentation system.

### qPCR

cDNAs were diluted 1:10 in ddH₂O. qPCR was conducted in a 96-well plate (Fisher Scientific, Cat. No. 44-833-54) using the QuantStudio 3 system (Thermo Fisher, Cat. No. A28567), with a final reaction volume of 20 µL per well (5 µL of diluted cDNA, 10 µL of SYBR Green master mix (Thermo Scientific, Cat. No. A46109), 0.5 µL of 10µM primer mix, and 4.5 µL of ddH₂O). Primer sequences for CNOT1, CNOT6, and CNOT9 are provided in **Supplemental Table S3**. Fast qPCR reactions began with an initial denaturation at 95°C for 2 minutes followed by 40 cycles of 95°C for 1s and 60°C for 30s. The C_T_/C_q_ values were then obtained using the Dx Real-Time PCR system software.

### Knockdown of CCR4-NOT subunits

DG75 or Mutu-BMRF1p-GFP cells (3-12 million) were initially transfected with 200 pmol of ON-TARGETplus non-targeting siRNAs (Dharmacon, Cat. ID. D-001810-01-20) or siRNAs targeting CNOT1 (Dharmacon, Cat. ID. J-015369-09-0010 or J-015369-11-0010), CNOT6 (Dharmacon, Cat. ID. J-019101-05-0010), or CNOT9 (Dharmacon, Cat. ID. J-019972-05-0010) by nucleofection (N=3 or N=4 per group). DG75 cells transfected with CNOT1 (or control) siRNAs were incubated for 48 hours. Due to time variation in knockdown efficiency for each on-target siRNA and for cell type, cells transfected with CNOT6 and CNOT9 siRNAs were incubated for 6 hours and 18 hours respectively in DG75, and 24 hours in Mutu-BMRF1p-GFP cells, post initial transfection. After these incubation times, cells were subjected to a second transfection as described below.

#### DG75 cells

Three million DG75 cells were co-transfected with the CMVp-GFP reporter (0.3 µg) and 3 µg of either a pLVX-BRRF1 expression vector or the empty vector control, pLVX-Cntl by nucleofection. Transfected cells were then incubated for 24 hours before collection for sorting of GFP+ cells by FACS.

#### Mutu-BMRF1p-GFP cells

Three million cells were co-transfected with 3 µg of either the SV40p-Zta or SV40p-Cntl plasmids with either 140 pmol of non-targeting or CNOT1-targeting siRNAs by nucleofection. The transfected cells were incubated for 24 hours before sorting of GFP+ cells by FACS.

### Generation of early gene and BRRF1 mutant expression plasmids

For each EBV early gene, the corresponding reading frame, with an in-frame C-terminal 3X FLAG sequence was synthesized by Synthego corporation and cloned into the lentiviral vector, pLVX-puro. BRRF1 mutants, BRRF1_C222A-I223A-Y224A-Y225A_ and BRRF1_Y224A_ were generated such that the codons for the indicated amino acid positions were substituted with an alanine codon.

### Co-immunoprecipitation

DG75 cells (24 million) were nucleofected with 24 µg of either the pLVX-Cntl plasmid or FLAG-tagged pLVX-BRRF1-WT, pLVX-BRRF1_Y224A_, or pLVX-BRRF1_C222A-I223A-Y224A-Y225A_. The cells were incubated for 24 hours and then resuspended in cold lysis buffer containing 1% Triton X-100 (Sigma Aldrich, Cat. No. X100-100ML), 50 mM HEPES (Boston BioProducts, Cat. No. BBH-74), 150 mM NaCl (Fisher Scientific, Cat. No. S271-1), 10% glycerol (Sigma Aldrich, Cat. No. G5516-500ML), 1.5 mM MgCl_2,_ (Thermo Scientific, Cat. No. AM9530G), 1.0 mM EGTA (Bioworld, Cat. No. 40520008-1), 1X cOmplete™ EDTA-free protease inhibitor (Sigma-Aldrich, Cat. No. 11836170001), and 1X Phosphatase Inhibitor (Roche, Cat. No. 049068450001) and incubated on ice for 15 minutes. Lysates were then spun down at 18,000xg for 15 minutes at 4°C. Anti-FLAG M2 magnetic beads (Sigma-Aldrich, M8823-1ML) were equilibrated in TBS (50 mM Tris-HCl, pH 7.4, Boston BioProducts, BBT-74-DR; 150 mM NaCl) and blocked with 1% BSA (Cell Signaling Technology, 9998S) in TBS for 45 minutes at room temperature with rotation prior to addition to cell lysates. Equilibrated beads were added to each cell lysate and then incubated overnight at 4°C on a rotator. Bead-lysate mixtures were subsequently washed twice with high-salt buffer (50 mM Tris-HCl, 1M NaCl, 1 mM EDTA (Thermo Scientific, Cat. No. AM9260G), 1%(v/v) NP-40 (Bioworld, Cat. No. 41430004-1), 0.1%(v/v) SDS (Invitrogen, Cat. No. 15553-027), 0.5%(w/v) Sodium deoxycholate (Sigma-Aldrich, Cat. No. D6750-25G)) and twice with low-salt buffer (500 mM Tris-HCl, 250 mM MgCl_2,_ 5%(v/v) Tween-20 (Sigma-Aldrich, Cat. No. P7949-500ML), 125 mM NaCl). Beads were then collected and resuspended in 1X Laemmli buffer (Sigma-Aldrich, 53401-1VL), boiled at 95°C for 10 minutes, and analyzed by Western blotting.

### Western blotting

Protein samples were loaded onto a 4-20% Mini-PROTEAN precast polyacrylamide gel (BioRad, Cat. No. 4568093) and subjected to electrophoresis at 130 V for 70 minutes. The proteins were then transferred to nitrocellulose membranes (BioRad, Cat. No. 1704159) using the Trans-Blot Turbo Transfer System (BioRad, Cat. No. 1704150). Following transfer, the membranes were blocked with TBS containing 0.1% Tween-20 and 5% BSA for 1 hour at room temperature. Primary antibodies, including anti-CNOT1 (Proteintech, Cat. No. 14276-1-AP), anti-CNOT7 (Proteintech, Cat. No. 14102-1-AP), anti-RQCD1 (CNOT9) (Proteintech, Cat. No. 22503-1-AP), and anti-FLAG (Sigma-Aldrich, Cat. No. F1804-200UG), were each diluted 1:1000 in 5% BSA TBST. Diluted primary antibodies were incubated with the membranes overnight at 4°C. The next day, the membranes were washed with TBST and incubated with a 1:15,000 dilution of secondary antibodies (Goat Anti-Rabbit 680RD (Licor, Cat. No. 926-68071), Goat Anti-Mouse 680RD (Licor, Cat. No. 926-32213), Goat Anti-Mouse 800CW (Licor, Cat. No. 926-32210), or Goat Anti-Rabbit 800CW (Licor, Cat. No. 926-33219)) in 5% BSA TBST at room temperature for 1 hour. After incubation, membranes were washed with TBST and imaged using an Odyssey CLx Imager (Licor).

### Proximity Ligation Assay combined with immunofluorescence assay

Mutu cells (4 million) were co-transfected with either the pLVX-Cntl or pLVX-BRRF1-FLAG plasmids (2 µg), and with either the SV40p-Zta or the SV40p-Cntl plasmids (5 µg) and incubated for 18 hours. Cells were then collected, washed, resuspended in PBS, and seeded onto slides. Cells were fixed with 4% paraformaldehyde (PFA) for 15 minutes at room temperature (RT), washed twice with PBS and permeabilized with 0.2% Triton X-100 for 5 minutes at RT. After permeabilization, cells were blocked using the blocking solution (provided in the probe kits) for 30 minutes at 37°C. Primary antibodies (anti-FLAG, anti-CNOT1, anti-CNOT9) were diluted 1:50 in antibody diluent (Sigma-Aldrich, Cat. No. DUO82008-8ML) and incubated with cells for 60 minutes at 37°C. Cells were then incubated with anti-mouse MINUS (Sigma-Aldrich, Cat. No. DUO92004-100RXN) and anti-rabbit PLUS (Sigma-Aldrich, Cat. No. DUO92002-100RXN) probes diluted 1:5 in antibody diluent for 1 hour at 37°C. Ligation and amplification stock solutions were prepared at a 1:5 dilution in ultrapure water. Cells were washed twice with 1X Wash Buffer A (Sigma-Aldrich, Cat. No. DUO82049-4L) for 5 minutes at RT and incubated with ligation solution containing ligase (1:40 dilution) for 30 minutes at 37°C. Cells were then washed twice with 1X Wash Buffer A for 2 minutes each and incubated with amplification solution containing polymerase (1:80 dilution) for 100 minutes at 37°C. Following amplification, cells were washed twice with 1X Wash Buffer B (Sigma-Aldrich, Cat. No. DUO82049-4L) for 10 minutes each at RT in the dark. Cells were then subjected to immunofluorescence staining and incubated with primary antibody, anti-BMRF1/anti-EaD (EMD Millipore, Cat. No. MAB8186) diluted 1:200 in antibody diluent for 1 hour at 37°C. Cells were washed twice with PBS, followed by incubation with goat anti-mouse IgG Alexa Fluor 488 secondary antibody (1:500 dilution in antibody diluent) for 30 minutes at 37°C. Finally, cells were washed twice with PBS and mounted using ProLong™ Glass Antifade Mountant with NucBlue™ Stain (Thermo Scientific, Cat. No. P36983)

### RNA sequencing

Total RNA samples with RNA Integrity Numbers (RIN) between 9-10 were used for RNA sequencing. RNA sequencing was performed by BGI and MedGenome using stranded PolyA library preparation kits and sequenced on the DNBseq platform with paired-end 150 bp reads, generating reads for approximately 100 million fragments per sample. Sequencing quality was assessed using fastp and fastqc prior to alignment. Reads were aligned to the human reference genome (GRCh38) combined with the Akata EBV genome using STAR v2.7.10b with the following parameters: **--chimSegmentMin 2, --outFilterMismatchNmax 3, --alignEndsType EndToEnd, --outSAMstrandField intronMotif, --alignSJDBoverhangMin 6, and --alignIntronMax 300000**. Resulting BAM files were analyzed using rMATS v4.3.0 (26), and downstream analyses were performed using Junction Counts and Exon Counts (JCEC) output files. RNA samples were additionally assessed for mycoplasma contamination, and strand specificity was evaluated using rseqc. Differential alternative splicing events (A3SS, A5SS, MXE, RI, and SE) were considered significant at FDR < 0.0005 with an inclusion level difference (IncDiff) threshold of 0 (Negative IncDiff < 0; Positive IncDiff > 0). The splicing events and their corresponding coordinates were visualized and validated using the Integrative Genomics Viewer (IGV).

#### Downstream analyses

##### Splicetools

For all modules in SpliceTools (27), an FDR threshold of < 0.0005 was used. For SEFractionExpressed module, a minimum TPM of 3 for either condition (control or test) was applied.

##### Conserved domain analyses

The amino acid sequences of all in-frame skipped exons were obtained using the SETranslateNMD module in Splicetools (27). These sequences were then queried against the NCBI Conserved Domain Database (https://www.ncbi.nlm.nih.gov/Structure/bwrpsb/bwrpsb.cgi) for domain homology analysis.

##### Gene ontology

Genes undergoing statistically significant increased exon skipping induced by reactivation models (FDR < 0.0000005) and by top five early genes (FDR < 0.0005) were extracted and subjected to pathway enrichment analysis using Enrichr (28–30).

### Image acquisition and processing

Images were acquired and edited using a Keyence BZ-X800 fluorescence microscope with z-stack (stepsize = 0.2 µm) acquisition under identical settings across all samples.

### 3D modeling, visualization, assessment of structural similarity and identification of interface residues

The interactions between: 1) BRRF1 and individual CCR4-NOT subunits, 2) BRRF1 mutants and CNOT9, 3) CDK2 and CCNE1/CCNA2, were predicted using AlphaFold3 (31), using default parameters. Three-dimensional structural models of BRRF1 mutants were generated accordingly. Predicted protein complexes, including protein-protein interaction, interacting interface residues, and structural alignments for similarity assessment were visualized and analyzed using PyMOL (version 3.1.6.1) (32). Structural similarity was assessed based on root mean square deviation (RMSD) values calculated in PyMOL (32), and interacting residues were defined based on a distance cutoff of <4 Å between protein chains using AlphaFold3-predicted structures (**Supplemental File S1**).

### Protein sequence/ID retrieval, protein sequence alignment and domain annotation

Canonical protein sequences and protein IDs for genes exhibiting significantly increased exon skipping and involved in innate immune signaling (**Supplemental Table S1**) were retrieved using Uniprot (33). Domain annotations were obtained using InterPro (34), with Pfam (35), powered by the HMMER3 package, used as the reference database for domain classification.

The protein sequences of BRRF1 and ORF49 were aligned using Clustal Omega (36), and protein sequence similarity searches were performed using BLASTp (NCBI) (37).

### Prediction of neopeptide antigenicity

The intrinsic antigenicity of each peptide sequence was predicted using IAPred, an open-source tool that applies a supervised machine-learning framework: a support vector machine (SVM) classifier trained on a curated set of pathogen-derived proteins to estimate intrinsic antigenicity from peptide sequences (38). All sequences were prepared in FASTA format. We used the authors’ reference implementation with a slightly customized pipeline, so the tabular output retained the input amino acid sequence together with score and category fields. Following IAPred’s default reporting, each scored peptide was classified as Low (intrinsic antigenicity score < −0.3), Moderate (−0.3 to 0.3), or High (> 0.3).

### Functional enrichment of highly antigenic neopeptides with STRING-db

We performed functional enrichment on the high intrinsic antigenicity gene set using STRING (v12, Homo sapiens, taxon 9606) (39). Enrichment was executed with a custom Python script that submits the gene list to the STRING-db REST API and saves the returned TSV results. Gene symbols were mapped to canonical identifiers within that pipeline and tested for over-representation across the enrichment category types returned by STRING. Terms with FDR < 0.05 in the STRING enrichment output were considered statistically enriched. In parallel, we used the STRING web server to explore protein-protein association networks, disease context, and enrichment results interactively.

### Functional analysis of genes with increased exon skipping in innate immune signaling

Statistically significant exon skipping (SE) events associated with genes involved in innate immune signaling were identified in the Mutu-Zta reactivation model (FDR < 0.0005, IncDiff < −0.1). These genes and their corresponding SE events were categorized into different axes of innate immune signaling. For each selected SE event, nonsense-mediated decay (NMD) (or frameshift) status was assessed using the SETranslateNMD module. Canonical protein sequences were obtained from UniProt, and protein domain annotations were retrieved from the Pfam database.

For NMD-targeted genes, frameshifts were assumed to initiate from the upstream exon end position (or upstream donor site), and potential domain loss was evaluated from this position to identify affected annotated domains. For non-NMD-targeted genes, overlap between skipped exon sequences and annotated protein domains was assessed.

## Results

### A multi-model framework for high-confidence analysis of lytic replication-induced host transcriptome remodeling

To define host transcriptome remodeling during EBV (and KSHV) lytic replication with a high degree of rigor and generalizability, we established a diverse panel of complementary reactivation systems spanning distinct viral, cellular, and induction contexts, paired with lytic cell purification strategies and high-depth sequencing. Specifically, we used five reactivation models including two EBV-positive Burkitt’s lymphoma cell lines, Akata and Mutu, induced through either B cell receptor (BCR) crosslinking or ectopic expression of the EBV transactivator, Zta (published previously) (3), the EBV-positive stomach cancer cell line, SNU719, transfected with Zta plus a second EBV transactivator, Rta, and the KSHV-positive TREX-BCBL1 cell line in which Rta expression can be induced through doxycycline treatment (40).

A central challenge in studying EBV and KSHV lytic replication is that only a subpopulation of cells enters the lytic cascade, with non-reactivated cells causing background signal that can obscure bona fide lytic-associated effects. To overcome this obstacle, we used Akata and Mutu cells carrying a BMRF1p-GFP lytic reporter and isolated GFP-positive cells in cultures treated with anti-Ig (to activate the BCR) and GFP-negative cells in mock-treated cultures (**Figure 1A**, *left panel*) (3). To ensure that any observed changes were the result of reactivation and not B cell receptor signaling, our third B cell model used Mutu cells co-transfected with a constitutive GFP expression vector and either a control or a Zta expression vector, with transfected cell populations being selected by FACS (**Figure 1A**, *right panel*). For the stomach cancer cell line, SNU719, cells carrying the BMRF1p-GFP lytic reporter were transfected with Zta and Rta to induce reactivation. GFP-positive (from Zta + Rta transfected) and GFP-negative (from control transfected) cells were then isolated 24 hours later. For all experiments, RNA was extracted from sorted cells and subjected to high-depth RNA sequencing of poly(A) RNAs. Finally, the KSHV BCBL1 cell line carrying an inducible Rta cassette (TREX-BCBL1) was induced with doxycycline for 24 hours and similarly subjected to high-depth RNA sequencing of poly(A) RNAs (40). Together, these models provide a rigorously controlled and broadly applicable framework for defining lytic replication-induced host transcriptome remodeling with high confidence and resolution.

**Figure 1.**
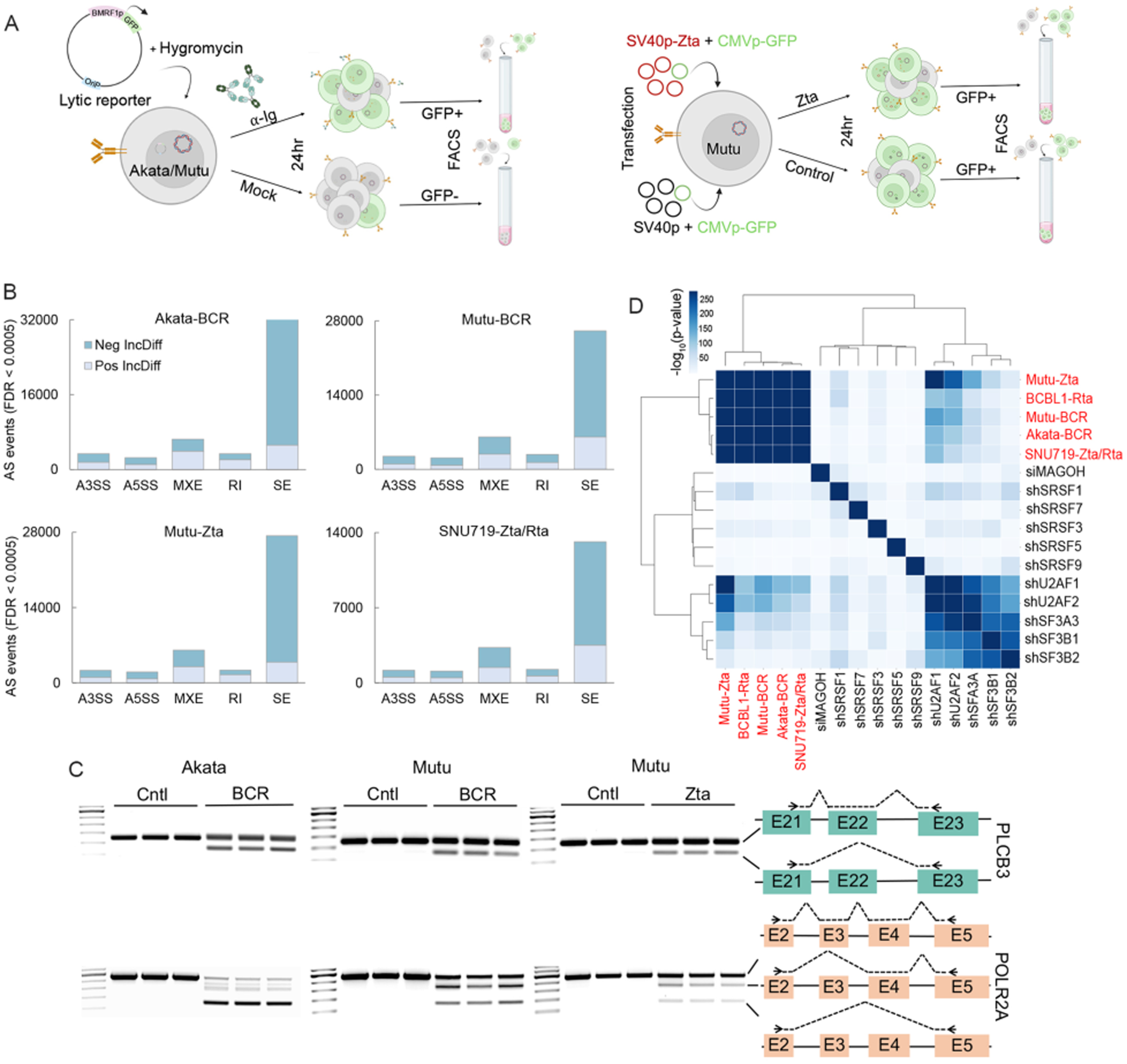
Extensive splicing alterations associated with gammaherpesvirus reactivation. **(A)** Reactivation models. (*left panel*) BCR crosslinking: The lytic reporter, pCEP4-BMRF1p-GFP was stably transfected into Akata and Mutu cells (and SNU719 cells -not shown). Akata-BMRF1p-GFP and Mutu-BMRF1p-GFP cells were treated with either anti-IgG (Akata) or anti-IgM (Mutu) antibodies (or mock-treated) for 24 hours. Induced GFP+ cells and latent (control) GFP-control cells were selected by FACS. *(right panel*) Ectopic Zta induction: Mutu cells were co-transfected with a CMVp-GFP reporter plasmid and either a SV40p-Cntl or a SV40p-Zta expression vector, GFP+ cells were collected 24 hours later, and RNA was isolated and subjected to RNA sequencing. This panel was created with the assistance of BioRender.com**. (B)** Splicing changes in EBV reactivation models. The number of alternative splicing (AS) events for each type was assessed using rMATS with a false discovery rate (FDR) < 0.0005. Inclusion level changes for each of the five AS types is represented as a positive inclusion difference (Pos-IncDiff, IncDiff > 0) or a negative inclusion difference (Neg-IncDiff, IncDiff < 0). **(C)** RT-PCR validation of induced SE events for the PLCB3 and POLR2A genes across all three BL EBV reactivation models. **(D)** Comparative analysis of common increased SE events (Neg-IncDiff) in EBV and KSHV reactivation models, and the knockdown of several core splicing factors using the SpliceCompare module (Splicetools, FDR < 0.0005).

### Lytic replication drives widespread and conserved host splicing disruption

To determine the extent to which lytic replication reshapes host pre-mRNA processing, we performed a global analysis of alternative splicing across EBV reactivation models. Splicing changes were identified using rMATS (replicate multivariate analysis of transcript splicing) (26) to identify statistically significant changes in 5’ splice site usage (A5SS), alternative 3’ splice site usage (A3SS), mutually exclusive exon usage (MXE), intron retention (retained intron (RI)), and exon skipping (skipped exon (SE)). Using a stringent false discovery rate (FDR) cutoff of 0.0005, extensive and reproducible splicing changes were observed across all three EBV Burkitt lymphoma models as well as the EBV gastric carcinoma reactivation model (SNU719), with especially high numbers of changes being observed for exon skipping (**Figure 1B**). We then validated a handful of induced exon skipping events, including exon skipping at the PLCB3 and POLR2A genes by RT-PCR (**Figure 1C**), confirming consistency across reactivation models.

Extending these analyses to KSHV, lytic replication was induced in the TREX-BCBL1 cell line through induction of Rta expression with doxycycline. In this model, we similarly observed substantial splicing alterations across all splicing types (**Supplemental Figure S1**), consistent with our recent report (41). Together, the observed splicing changes in all four EBV reactivation models tested and KSHV-infected BCBL1 cells support the contention that extensive alternative splicing is a conserved feature of reactivation in human gammaherpesviruses.

Given the broad splicing disruption observed across EBV and KSHV models, we investigated whether these changes converge on common mechanistic features by assessing the overlap of common increased exon skipping events (Negative IncDiff). Using the SpliceCompare module of our published software suite, SpliceTools (27), which determines the statistical significance of overlaps, we observed a strong clustering of all EBV and KSHV reactivation models for increased exon skipping events (**Figure 1D**). We also observed a smaller but significant overlap in splicing changes between our reactivation models and the knockdown of several core U2 splice-acceptor assembly factors (1), with the most pronounced overlap being with the knockdown of U2AF1 and U2AF2 (**Figure 1D**).

Altogether, these findings indicate that EBV and KSHV reactivation causes substantial splicing alterations and that the SE changes across reactivation models share mechanistic similarities which may occur, in part, through disruption of splice-acceptor activity.

### Lytic replication drives transcriptome-wide disruption of host splicing

Among the five AS types, exon skipping was the most prevalent in each of the EBV and KSHV models, accounting for up to 68% of all statistically significant AS events (**Supplemental Figure S1**). Increased exon skipping predominated over exon inclusion (Positive IncDiff) (**Figure 2A**), consistent with reactivation-associated loss of normal splicing fidelity. The degree of exon skipping caused by reactivation was extensive, outweighing the number of statistically significant increased exon skipping events observed by the knockdown of any of 186 RNA-binding proteins (RBPs) and splicing factors (1,27), including the key splicing factors, U2AF1 and U2AF2 (**Figure 2A**). Further evidence for the extent of splicing disruption was the finding that while single exon skipping was predominant, exon skipping events frequently involved the skipping of multiple exons (**Figure 2B**). Together, these findings highlight the substantial magnitude of reactivation-induced exon skipping and suggest that lytic replication broadly compromises the integrity of host mRNA processing.

**Figure 2.**
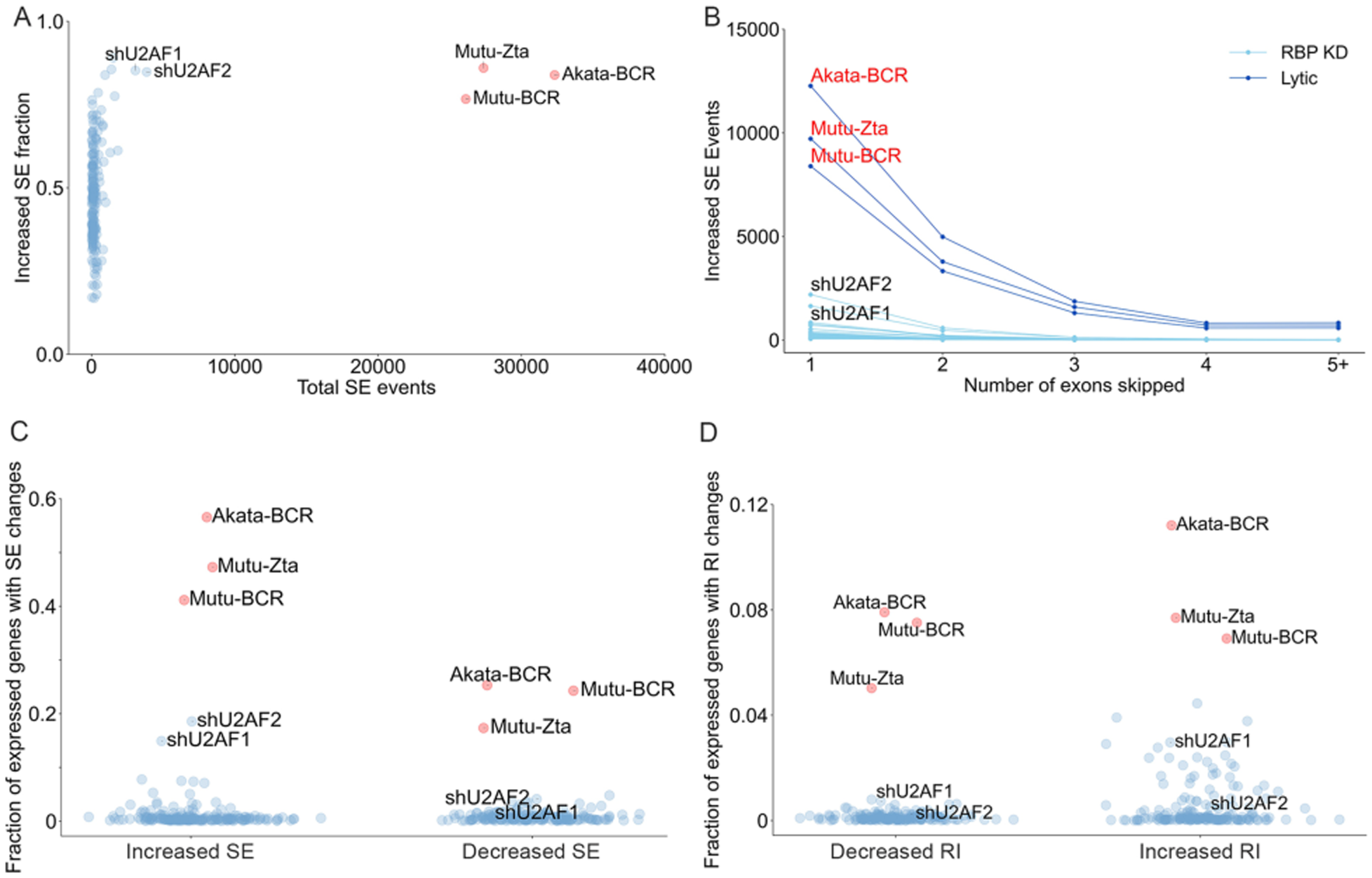
Footprint of alternative splicing across the cell transcriptome. **(A)** Reactivation induces extensive exon skipping. Total significant (increased + decreased) exon skipping events (x-axis) and fractions of increased exon skipping (y-axis) events comparing three EBV reactivation models to the knockdown of 186 RNA-binding proteins (RBPs). SE analyses were assessed using rMATS with FDR < 0.0005. The dataset of 186 RBP knockdowns in HepG2 cell lines were obtained from NCBI Sequence Read Archive or ENCODE project (1) (See **Supplemental File S2** for accession numbers). **(B)** Degree of multiple exon skipping. The number of significant increased events with each of one or more exons skipped across three EBV reactivation models and RBP knockdowns was assessed using the SENumberSkipped module (SpliceTools, FDR < 0.0005). For the RBP knockdown dataset, only the knockdowns with statistically significant increased SE events of > 100 were selected. Penetrance of exon skipping (SE) **(C)** and intron retention (RI) **(D)** changes across the expressed transcriptome in reactivation models and RBP knockdown datasets. The fraction of expressed genes with increased and decreased SE or RI was assessed using SEFractionExpressed or RIFractionExpressed module (Splicetools, FDR < 0.0005) respectively, with a minimum TPM of 3 in either control or test condition.

To determine how deeply splicing disruption extends into the host transcriptome, we next quantified the fraction of expressed genes affected by increased exon skipping using the SpliceTools FractionExpressed module (27). Analyzing all genes expressed with a minimum of 3 TPM (transcripts per million), up to 56.6% exhibited one or more statistically significant (FDR threshold of 0.0005) increased exon skipping event during reactivation (**Figure 2C**), substantially exceeding the fraction of expressed genes showing increased exon skipping for any RBPs knockdown. This degree of penetrance reveals that reactivation-associated splicing disruption is not limited to selected targets, but instead extends broadly across the expressed host transcriptome, with likely widespread functional consequences.

Because intron retention can also have a significant impact on transcript output and gene function, we next examined the pervasiveness of significant changes in this class of splicing alterations. This analysis showed that intron retention changes accounted for 5.9-9.0% of all statistically significant AS events among our four EBV reactivation models (**Supplemental Figure S1**). Assessing the penetrance of these changes across the expressed host transcriptome, we found that 6.9-11.2% of expressed genes displayed statistically significant changes in increased intron retention (Positive IncDiff) (**Figure 2D**). Considering exon skipping and intron retention alone, lytic reactivation alters RNA processing across a substantial portion of the host transcriptome, with likely widespread consequences for cellular gene output.

### EBV-induced splicing changes are predominantly disruptive and functionally deleterious

The analyses presented above support a model in which lytic replication broadly disrupts normal splicing function rather than selectively engaging physiological splicing regulatory programs to rewire host pathways. To further test whether lytic replication causes splicing disruption rather than regulated splicing reprogramming, we quantified the fraction of induced splicing events that are not represented in existing transcript annotations. As a proxy for disruption, we reasoned that splicing events associated with normal cellular regulatory programs are more likely to have been previously observed and incorporated into Ensembl annotations, whereas disruptive splicing should generate a higher fraction of unannotated events. Using the FractionUnannotated module of SpliceTools (27), we found that 70.5-75.5% of all statistically significant increased SE events were not previously annotated (**Figure 3A**). Moreover, these values exceeded those observed following knockdown of core splicing factors such as MAGOH, U2AF1, and U2AF2. These data support the conclusion that EBV reactivation does not primarily remodel established splicing programs but instead exerts a broadly disruptive effect on host splicing fidelity.

**Figure 3.**
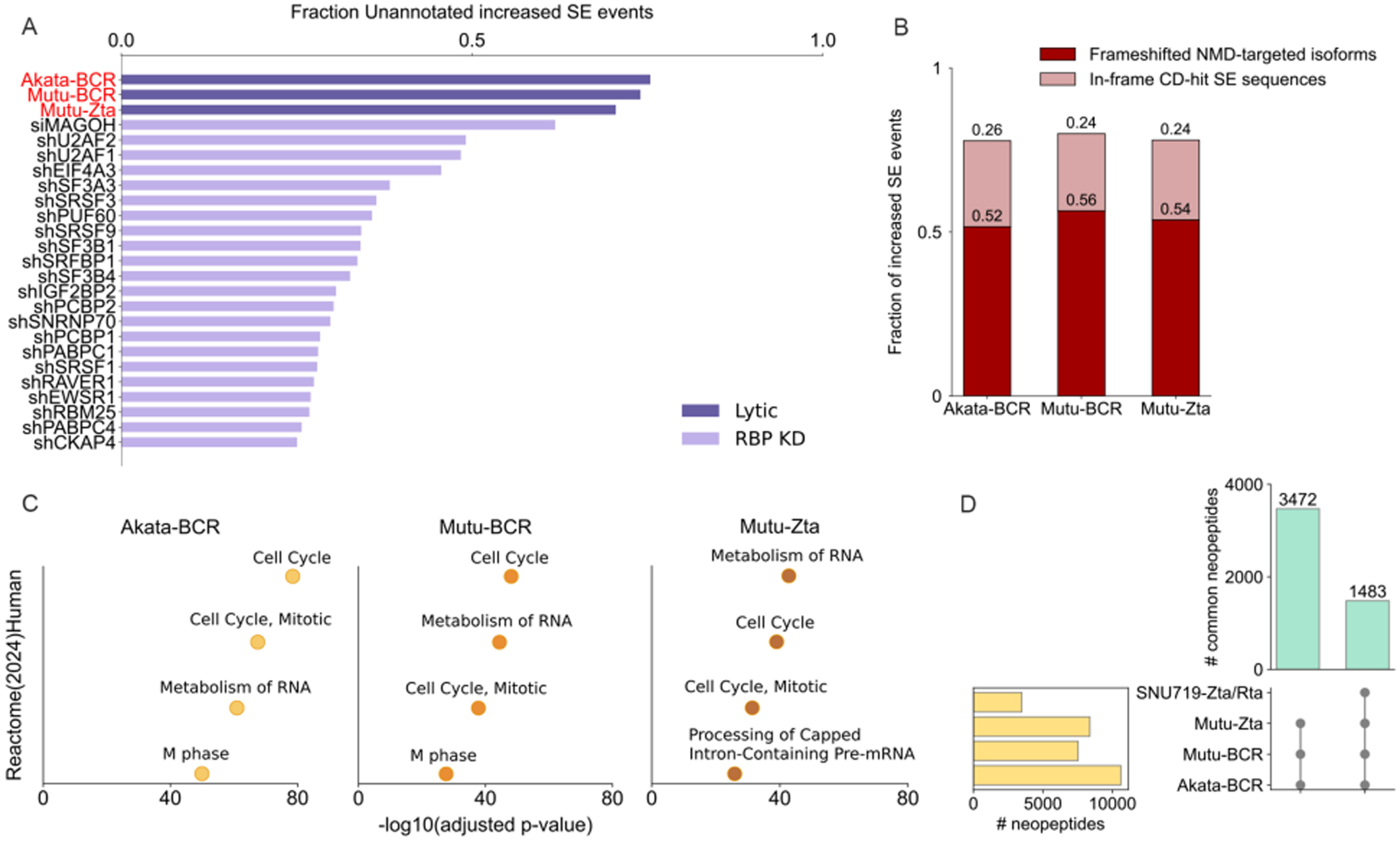
Disruption of cell splicing and functional impact prediction. **(A)** Fraction of unannotated SE events. The fraction of significant increased SE events that were not previously annotated was compared across three EBV reactivation models and the knockdown of RBPs using SEFractionUnannotated module (Splicetools, FDR < 0.0005). For the RBP knockdown datasets, only top 25 of RBPs that showed the greatest number of significant increased SE events are displayed. **(B)** Functional impact of increased exon skipping on protein function. Nonsense-mediated decay (NMD) predictions for frameshifted SE transcripts in the three EBV reactivation models were generated using SETranslateNMD module (Splicetools, FDR < 0.0005). For in-frame SE events, the resulting amino acid sequences of all in-frame skipped exons from SETranslateNMD analysis were queried against the NCBI Conserved Domain Search Database (https://www.ncbi.nlm.nih.gov/Structure/bwrpsb/bwrpsb.cgi) to assess the presence of conserved domains. **(C)** Genes with significantly increased SE events from EBV reactivation models were identified using an FDR threshold of < 0.0000005. These genes were then subjected to pathway enrichment analysis using Enrichr. Displayed are the top four statistically significant pathways enriched from the Reactome Human (2024) database. (**D**) Frameshift-derived neopeptides during EBV reactivation. The number of neopeptides in each EBV reactivation model, as well as shared neopeptides among EBV-positive B cell reactivation models or between B and epithelial reactivation models, were assessed using the SETranslateNMD module of SpliceTools (FDR < 0.0005).

To assess the likely functional consequences of increased exon skipping during reactivation, we next asked how often these events are predicted to impair productive host gene output. Using the SETranslateNMD module of SpliceTools (27), we found that 52-56% of increased SE isoforms were predicted to introduce frameshifts leading to nonsense-mediated RNA decay (NMD) (**Figure 3B**), indicating a substantial negative impact on host gene expression and/or the production of non-functional protein products due to frameshifts. For the remaining 44-48% of increased SE isoforms that preserved the reading frame, we evaluated whether the skipped exon encoded sequences correspond to known conserved protein domains. Using the amino acid sequences of each in-frame skipped exon and the NCBI batch Conserved Domain Search program (https://www.ncbi.nlm.nih.gov/Structure/bwrpsb/bwrpsb.cgi), we found that up to 26% of increased SE events removed an exon with homology to one or more conserved domains (**Figure 3B**). Together, these findings indicate that EBV-induced exon skipping is predicted to broadly compromise host proteome integrity by reducing transcript abundance through NMD and/or generating truncated proteins, and through generating proteins lacking functionally important domains.

### Predicted impact on cell signaling and immunogenicity

To investigate pathways predicted to be altered by increased exon skipping, we used Enrichr (28–30) to identify over-represented pathways for genes with statistically significant increased exon skipping (using an FDR cutoff of 0.0000005). The most significantly enriched pathways identified in this analysis were cell cycle and RNA metabolism (Reactome human database) (**Figure 3C**). The enrichment in RNA metabolism pathways, which includes splicing factors, indicates the likely presence of a self-reinforcing feedback loop in which EBV induced splicing disruption impacts the processing of transcripts encoding RNA splicing factors, which in turn, further reinforces disruption of these pathways.

Because EBV is associated with a variety of autoimmune diseases (42–44), we investigated whether the extensive splicing alterations reshape the host epitope landscape through translation of frameshifted mRNAs. Using SETranslateNMD module of SpliceTools (27), we found that EBV reactivation generates a large number of host neopeptides, ranging from 3480 (SNU719-Zta/Rta) up to 10,612 neopeptides (Akata-BCR) (**Figure 3D, Supplemental File S3**). Computational prediction of peptide antigenicity of events shared across models identified hundreds of neo-epitopes with moderate or high predicted antigenicity (**Supplemental File S3 and Figure S2**). Assessing tissue enrichment for genes associated with neo-peptides with high predicted antigenicity identified reproductive tissues and brain and nervous system (**Supplemental Figure S3**), the latter two of which are notable due to EBV’s association with MS. Notably, these host neo-epitopes could have an impact on host protein recognition beyond the neo-epitope region through epitope spreading to previously tolerized regions of host proteins. This is a concept that warrants further investigation with respect to EBV-associated autoimmune diseases.

Previous studies have provided evidence that EBV lytic replication leads to a block in cell cycle progression at the G1-S border (45–50). This pseudo-S-phase environment is thought to favor viral DNA replication by ensuring high levels of nucleotide pools needed for DNA replication without competition for these resources by cell DNA replication. Although the arrest at the G1-S border is thought to occur due to the activation of a DNA-damage response that is induced by viral lytic replication, we identified exon skipping for cell cycle genes that are key regulators of the G1 to S phase transition. Specifically, we identified a skipped exon isoform of CCNE1 (Cyclin E1), which is important for binding and activating CDK2 to facilitate the G1/S phase transition (51–53). In this isoform, exon 8 is consistently skipped across reactivation models (**Figure 4A**). This SE event generates an in-frame transcript in which the deleted region overlaps partially with the C-terminal cyclin box domain. This domain in Cyclin E exhibits significant differences in conformation and sequence compared to cyclin A, enabling Cyclin E to have additional interactions with pCDK2 and thereby stabilizing the cyclin E1-CDK2 interface (**Supplemental Figure S4 and Figure 4A**) (54). Moreover, this region has been shown to contain the principal contact residues required for interaction with phosphorylated CDK2 (pCDK2) in the pCDK2/cyclin E1 complex (54). Using AlphaFold3 to predict the interaction between canonical CCNE1 and CDK2 (UniProt (CCNE1: P24864; CDK2: P24941)) isoforms, we found a high-confidence interaction between full length CCNE1 and CDK2 (**Figure 4A**). Using the skipped exon isoform of CCNE1 lacking the exon-skipped sequence, however, showed a substantially lowered interaction score (**Figure 4A**).

**Figure 4.**
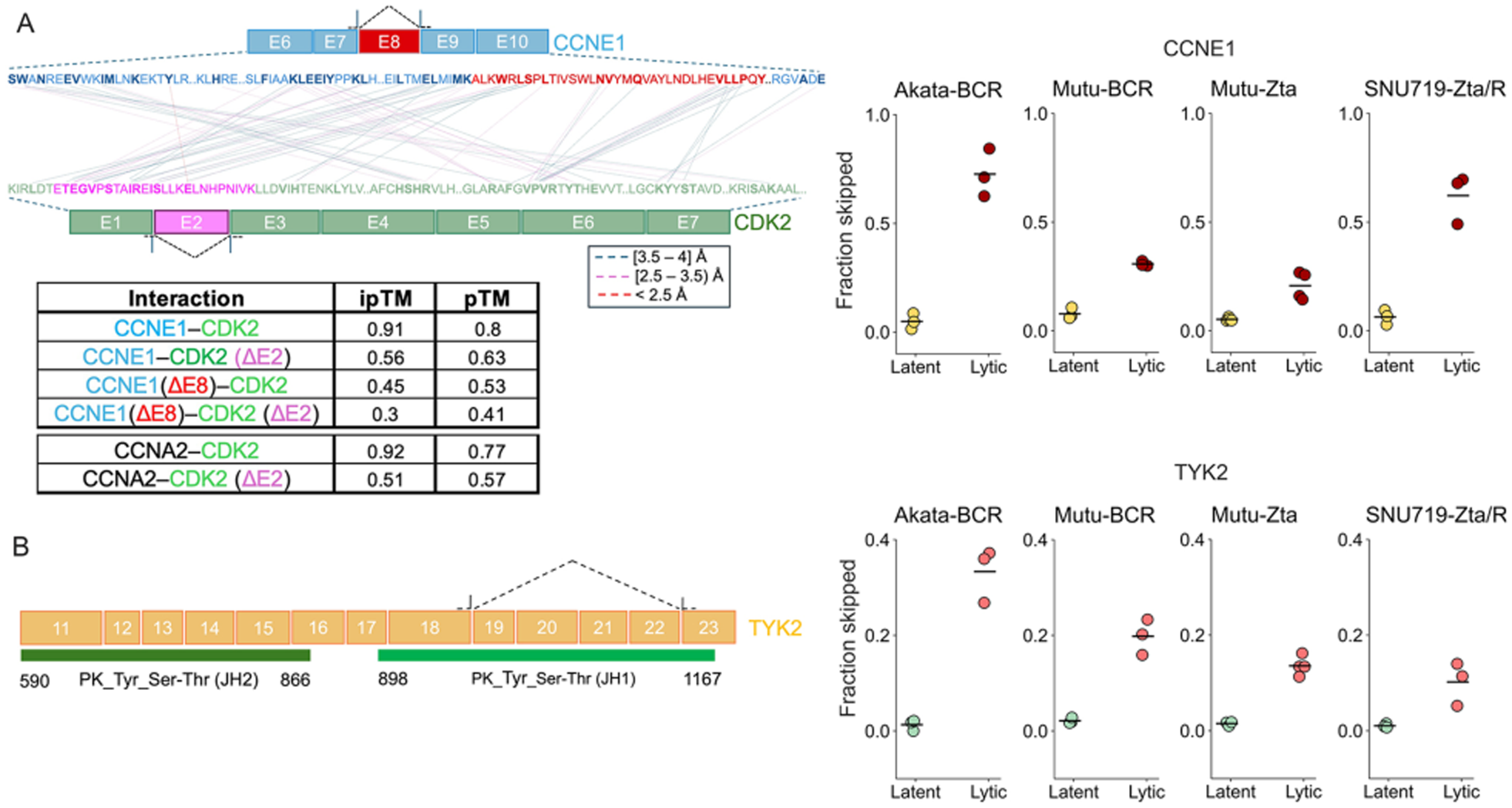
Predicted impact of increased exon skipping on host pathways. (**A**) Cell cycle pathway. Exon skipping of CCNE1 exon 8 and CDK2 exon 2 were identified as significantly increased from rMATS (FDR < 0.0005). CCNE1 and CDK2 directly interact to regulate the G1/S transition. (*Left*) The interactions between CCNE1 isoforms and CDK2 isoforms, as well as between CCNA2 and CDK2 isoforms were predicted using AlphaFold3, with predicted interaction scores shown in the table. Interface residues between CCNE1 and CDK2 were identified using PyMOL with a distance cutoff of <4 Å and categorized as very proximal (<2.5 Å, red), moderate (2.5-3.5 Å, pink), or distant (3.5-4 Å, blue) (see **Supplemental File S1** for interface residues). These analyses indicate that regions outside exon 8 of CCNE1, as well as regions outside exon 2 of CDK2, also contribute to the interaction. (*Right*) The fraction of exon skipping was calculated based on junction read counts, where skipped junctions (CCNE1: 29821817-29822239; CDK2: 55967124-55968048) and individual junctions (CCNE1: 29821817-29821995 and 29822130-29822239; CDK2: 55967124-55967856 and 55967934-55968048) were used to compute fraction skipped across EBV reactivation models (skipped junction counts divided by the sum of skipped junction counts and the average of individual junction counts). Each dot represents an individual replicate, and horizontal bars indicate the mean. (**B**) JAK-STAT signaling pathway. (*Left*) Representation of the exon-skipping event in TYK2, in which exons 19-22 are skipped (FDR < 0.0005). The kinase (JH1, aa 898-1167) and pseudokinase (JH2, aa 590-866) domains are annotated based on the Pfam database integrated into InterPro. (*Right*) The fraction skipped was calculated similarly to the CCNE1 and CDK2 fraction skip in (**A**).

We also observed that CDK2 undergoes statistically significant increased exon skipping of exon 2 across our reactivation models (**Figure 4A).** The second exon of CDK2 contains the C-(PSTAIRE) helix which is classically considered important for binding to cyclin E and cyclin A (**Figure 4A**) (52,54). The relevance of this region in binding to cyclin E is supported by AlphaFold3 prediction where a reduced interaction between wild-type CCNE1 and exon 2-deleted CDK2 is predicted (**Figure 4A**). Further, modeling the interaction between the exon 2-deleted CDK2 and the exon 8-deleted CCNE1 shows a nearly abolished interaction score (**Figure 4A**). Finally, since CDK2 also binds cyclin A to facilitate progression through S-phase, we assessed the impact of exon 2 deletion on this interaction. Whereas AlphaFold3 predicts a strong interaction between CDK2 and CCNA2 (CCNA2: P20248), binding of exon 2-deleted CDK2 and CCNA2 is similarly diminished (**Figure 4A**).

Together, with these interactions being critical for the G1 to S phase transition (cyclin E) and progression through S-phase (cyclin A), splicing disruption may serve as a layered mechanism to enforce a block to entry into and through S-phase, thereby reducing competition between viral and host DNA replication for nucleotide precursors.

In response to reactivation, specific cell signaling pathways are also activated (55–57). While signaling pathways required for viral persistence and cell survival during latency are already active and supported by pre-existing pools of functional proteins, the signaling pathways that are activated only in response to reactivation may be more vulnerable to disruption by these non-functional protein products of skipped exon isoforms. As a result, extensive host splicing disruption is likely to have a greater negative impact on host gene-expression pathways that are newly activated in response to lytic replication, the most salient of which would be innate immune signaling. Strikingly, despite the extensive nature of complex and antisense viral transcription as well as extensive host transcription remodeling, we observed only minimal expression of interferon response genes (3,58). Other mechanisms have previously been identified that help explain the blunting of interferon signaling during reactivation but alternative splicing of either downstream interferon response genes or regulators of IFN signaling could also contribute to suppression of interferon signaling. As shown in **Supplemental Table S1**, increased exon skipping is observed in a wide array of genes involved in interferon responses ranging from sensors to an array of signal transduction effectors including IKBKB, CHUK, NFKB1, TBK1, TANK, IRF1, IRF3, JAK1, TYK2, STAT2, and STAT5B. For example, reactivation promotes exon skipping of exons 19 through 22 in TYK2, a tyrosine kinase involved in JAK-STAT cytokine signaling and interferon responses (**Figure 4B**). This results in a frameshifted isoform that is not predicted to undergo NMD but encodes a protein lacking coding sequences within exons 19-22, including the kinase (JH1) domain.

Previous studies have shown that deletion of either the JH1 kinase domain or the JH2 pseudokinase domain, or both, abolishes autophosphorylation and phosphorylation of exogenous substrates (59). Consistent with this, kinase-deficient forms of JAK proteins (including TYK2) have been shown to function in a dominant-negative manner to repress IFN-α signaling (60). These observations suggest that EBV-induced splicing disruption can generate dominant-negative or nonfunctional proteins that impair antiviral signaling.

Altogether, the extensive nature of splicing disruption during lytic replication may not only lead to global decreases in the burden on the translation machinery but it may also lead to the generation of non-functional or dominant-negative proteins that support pathway disruptions that are critical for successful viral replication, such as cell cycle and innate immune signaling.

### Early gene expression underlies splicing disruption

To begin defining the mechanism underlying splicing disruption, we first tested whether this phenotype depends on viral DNA replication and the late phase of the lytic cycle. Mutu cells were transfected with a Zta expression vector to induce reactivation and cultured in the absence or presence of the viral DNA replication inhibitor, phosphonoacetic acid (PAA). As expected, PAA did not impair early gene expression but suppressed the expression of leaky late and late genes, consistent with effective inhibition of viral DNA replication (**Figure 5A**). Despite this block in viral DNA replication, PAA had little effect on the number of splicing changes across all five-alternative splicing (AS) classes (**Figure 5B**) and the fraction of increased exon skipping events that were not previously annotated (**Figure 5C**). Further, comparison of exon skipping amplitudes on an event-by-event basis showed that PAA had minimal impact on the magnitude of exon skipping (**Figure 5D**). Together, these findings indicated that splicing disruption during reactivation occurs largely independently of viral DNA replication and late gene expression, implicating one or more early viral gene products as candidate mediators.

**Figure 5.**
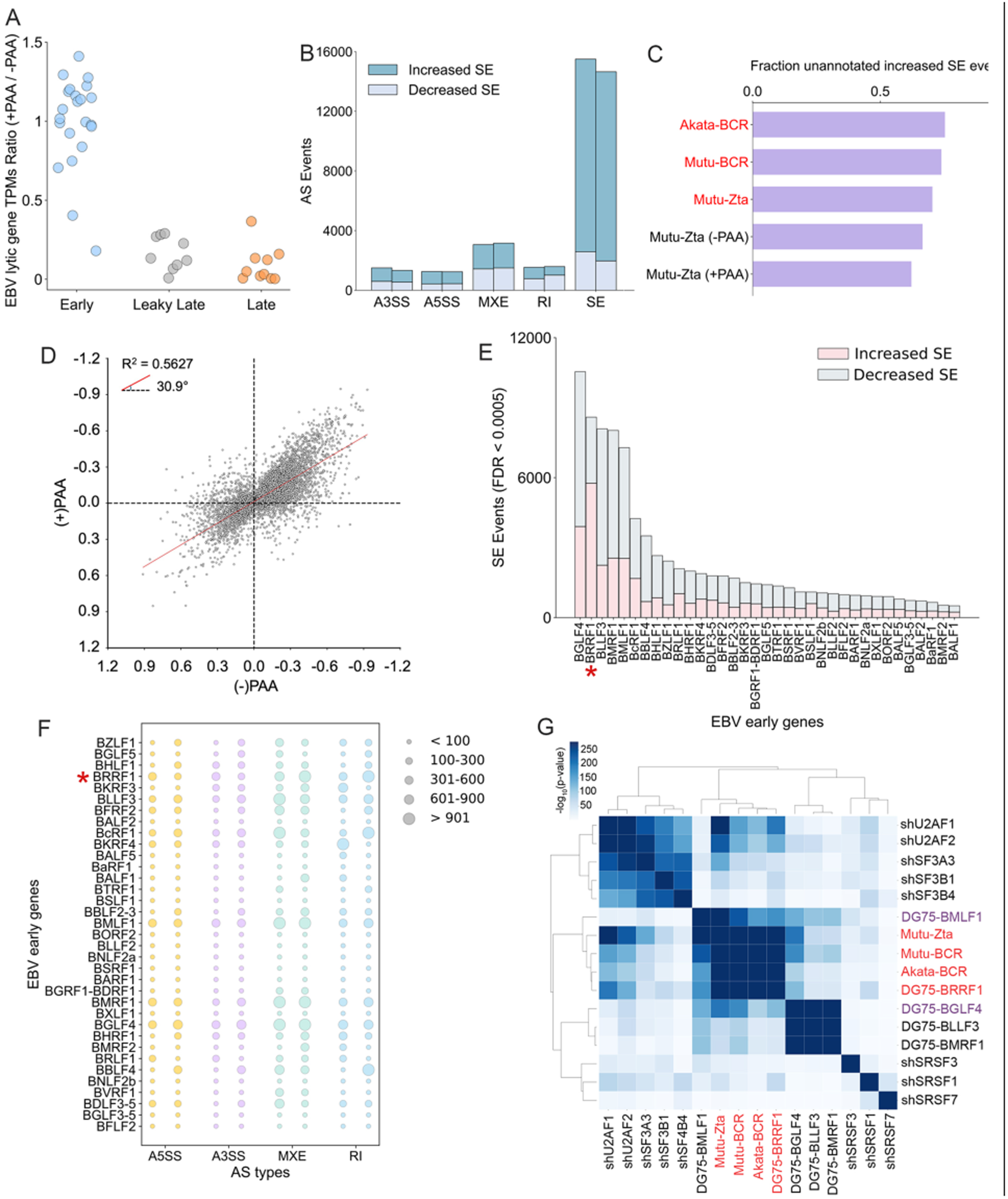
EBV splicing disruption is an early event with BRRF1 emerging as a prominent driver. **(A)** Lytic gene expression in the presence and absence of viral DNA replication inhibitor, phosphonoacetic acid (PAA). Mutu cells were reactivated via ectopic Zta expression with PAA (+PAA) or without PAA(-PAA) (N=4). The plot shows the ratio of average TPM values for the +PAA condition over the -PAA condition for each lytic gene. **(B)** Differential alternative splicing in Zta-reactivated Mutu cells with and without PAA. The number of statistically significant events for A5SS, A3SS, MXE, RI, and SE were assessed using rMATS (FDR < 0.0005). **(C)** The fraction of significant increased SE events that were unannotated across three EBV B cell reactivation models and PAA-treated/-untreated conditions (FDR < 0.0005). **(D)** Impact of PAA on splicing change magnitude. The Inclusion Difference (IncDiff) values of all statistically significant increased SE events (IncDiff_threshold_ = 0 and FDR < 0.0005) in the -PAA condition were compared with the corresponding IncDiff values of matching SE events in the +PAA condition. The coefficient of determination (R²) was determined and the angle of the regression line with x-axis was calculated from the arctangent of its slope. (**E, F**) The number of statistically significant SE **(E)** or A5SS, A3SS, MXE, and RI **(F)** events were assessed using rMATS (IncDiff_threshold_ = 0 and FDR < 0.0005). **(G)** Comparative analysis of common increased SE events between EBV reactivation models, the top 5 early genes with the greatest impact on exon skipping, and knockdowns of several core splicing factors. The analysis was performed using the SpliceCompare module (Splicetools, FDR < 0.0005).

### BRRF1 recapitulates features of reactivation-associated splicing disruption

To identify early genes capable of perturbing splicing, we generated expression vectors for EBV early genes and co-transfected each of them, together with a GFP expression vector, into the EBV-negative Burkitt lymphoma cell line, DG75. GFP-positive cells were isolated by FACS 24 hours later, and poly(A)+ RNA sequencing was performed on RNA extracted from the sorted cells (**Supplemental Figure S5A**).

Assessing splicing changes caused by individual EBV early genes using an FDR cutoff of 0.0005, we found that BGLF4, BRRF1, BLLF3, BMRF1, and BMLF1 exerted the greatest effects on splicing (**Figure 5E**). Among these candidates, BRRF1 was distinguished by the extent to which its splicing phenotype resembled the disruptive pattern observed during reactivation. Specifically, BRRF1 expression showed (1) a predominance of induced exon skipping over exon inclusion (**Figure 5E**), (2) the highest fraction of expressed genes with increased exon skipping events (**Supplemental Figure S5B**), and (3) the highest fraction of induced exon skipping events at previously unannotated loci (**Supplemental Figure S5C**). In addition, BRRF1 expression was associated with one of the highest numbers of significant increased intron retention events (**Figure 5F**).

To more directly assess the potential contribution of BRRF1 relative to BGLF4, BLLF3, BMRF1, and BMLF1 to splicing disruption during reactivation, we next examined the statistical significance of overlap between increased exon-skipping events induced by each early gene and those observed in reactivation models. While BLLF3, BMRF1, and BGLF4 showed significant overlap amongst themselves, BRRF1 clustered separately and showed the highest degree of overlap with all three reactivation models, with BMLF1 and BGLF4 also overlapping the reactivation models to a lesser extent (**Figure 5G**). These findings suggest that BRRF1 expression recapitulates a substantial component of the splicing disruption program observed during reactivation, therefore identifying BRRF1 as a strong candidate driver of this phenotype.

### The CCR4-NOT complex contributes to BRRF1-mediated splicing disruption

To investigate how BRRF1 disrupts splicing, we first identified candidate cellular interaction partners of BRRF1 using the EBV protein interactome resource generated by the Gewurz laboratory (61). Strikingly, BRRF1 was found to associate with multiple subunits of the CCR4-NOT complex, including CNOT1-3, CNOT6/6L, CNOT7, CNOT9 (RQCD1), and CNOT10 (61). We validated these interactions by co-immunoprecipitation in DG75 cells transfected with a FLAG-tagged BRRF1 expression vector, confirming association of BRRF1 with the core scaffold subunit CNOT1 as well as the accessory factors CNOT7 and CNOT9 (**Figure 6A**).

**Figure 6.**
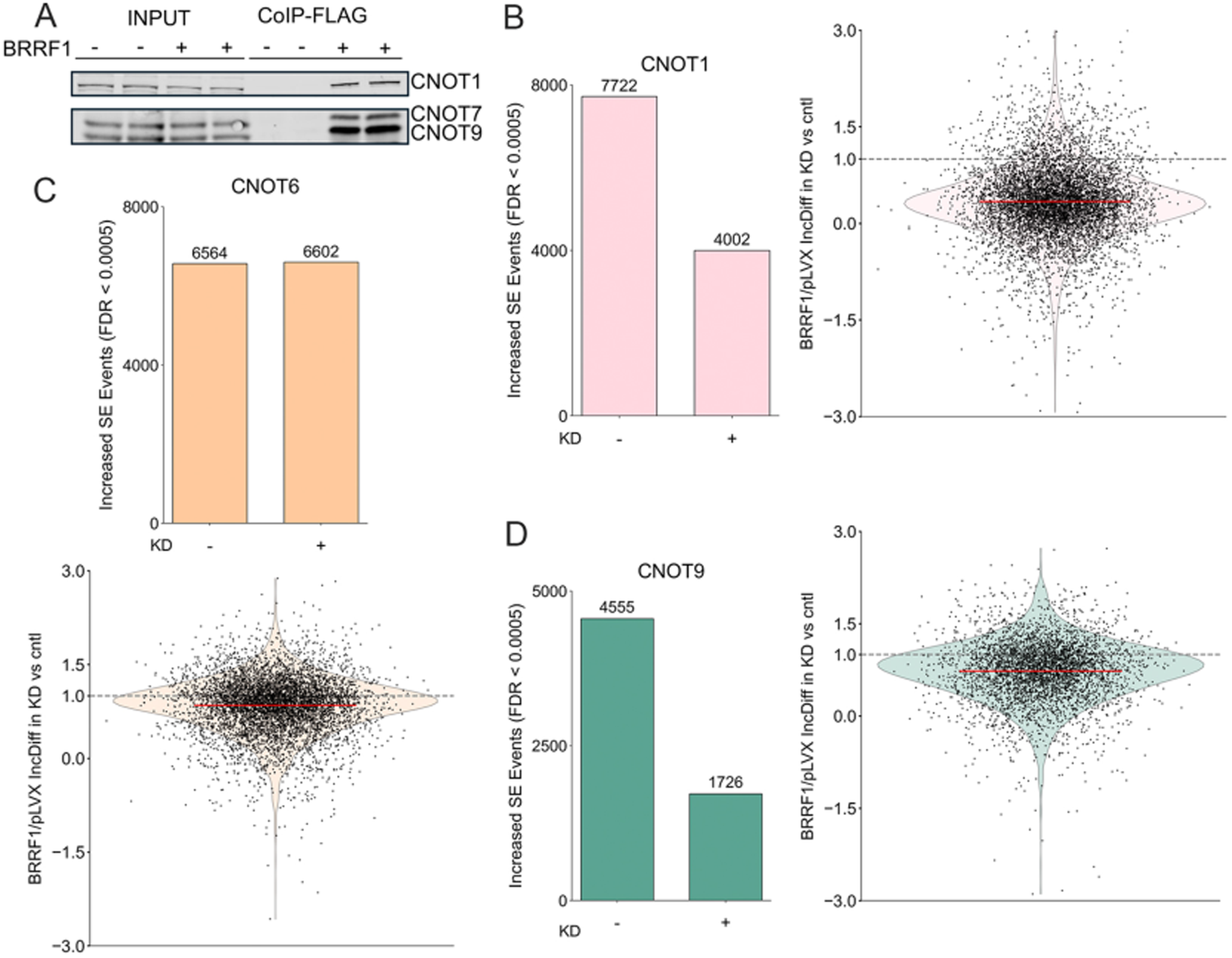
BRRF1 induces splicing changes through the CCR4-NOT complex. **(A)** Validation of BRRF1 and CCR4-NOT complex interaction via co-immunoprecipitation (Co-IP). DG75 cells were transfected with pLVX-puro plasmids expressing FLAG-tagged BRRF1 or empty-FLAG control. Anti-FLAG beads were used for the immunoprecipitation followed by western blot to detect CCR4-NOT subunits. **(B-D)** Knockdown of CCR4-NOT subunits with BRRF1 expression in DG75 cells. rMATS was used to assess changes in the number of significant increased SE events and the amplitude of SE changes in control knockdown (mock; siCntl-BRRF1 vs. siCntl-plVX; see **Supplemental Figure S6**) and CNOT knockdown (siCNOT-BRRF1 vs. siCNOT-pLVX; see **Supplemental Figure S6**) conditions (FDR < 0.0005). Number of statistically significant increased SE events following knockdown of CNOT1 **(B),** CNOT6 **(C)**, or CNOT9 **(D)** compared to mock was shown as bar plots. To assess the amplitude of BRRF1-induced SE changes upon CCR4-NOT subunit knockdown, Inclusion Difference (IncDiff) values for all significantly increased SE events identified under mock (siCntl-BRRF1 vs. siCntl-plVX) were extracted. Corresponding IncDiff values for the same SE events were obtained from each knockdown condition (siCNOT-BRRF1 vs. siCNOT-pLVX). Violin plots show the ratio of IncDiff values in CNOT1 (**B**), CNOT6 (**C**), or CNOT9 (**D**) knockdown relative to mock. A ratio < 1 indicates a reduction in the amplitude of BRRF1-induced SE events upon knockdown, whereas a ratio = 1 indicates no change.

Although the CCR4-NOT complex is best known for its predominantly cytoplasmic role in mRNA deadenylation, previous studies have also reported nuclear localization and nuclear functions for this complex (62–65). Because splicing is primarily a nuclear process, we examined the localization of CNOT1 and CNOT9 with FLAG-tagged BRRF1 under reactivated conditions. Using proximity ligation assays, we found that BRRF1 is in close proximity with both CNOT1 and CNOT9 in the nucleus of lytic cells (**Figure 7A and 7B**). Together, these findings support the possibility that BRRF1 perturbs splicing through its interaction with the CCR4-NOT complex in the nucleus.

**Figure 7.**
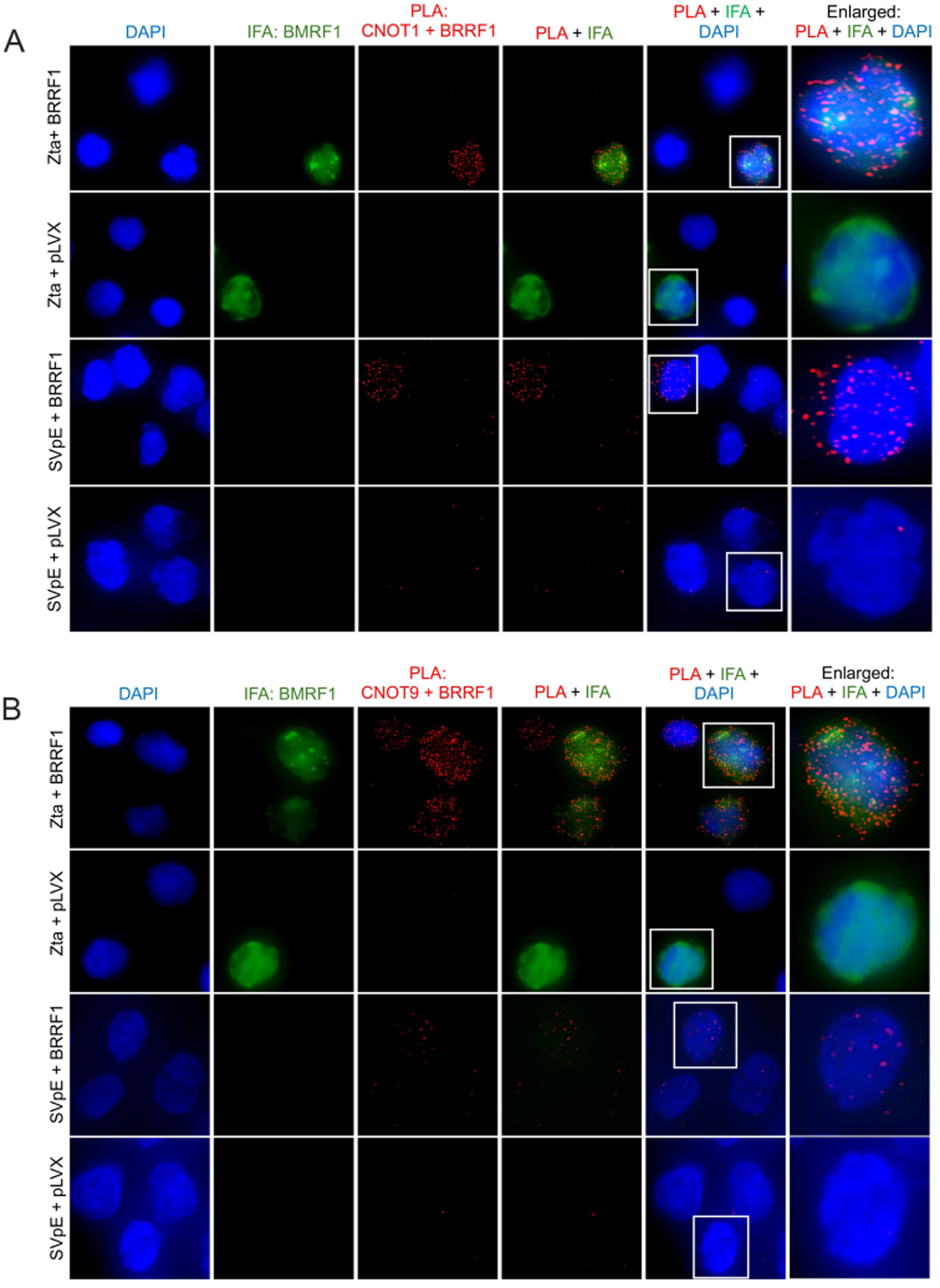
Nuclear (and cytoplasmic) interactions between FLAG-tagged BRRF1 and CNOT1 and CNOT9 in latent and reactivated cells. Mutu cells were co-transfected with either SV40p-Cntl (latent) or SV40p-Zta (reactivated) plasmids, together with either pLVX-Cntl or pLVX-BRRF1-FLAG and cultured for 18 hours. The subcellular localization of the interaction between FLAG-tagged BRRF1 and (**A**) CNOT1 and (**B**) CNOT9 was assessed by Proximity Ligation Assay (PLA). PLA was combined with IF, in which BMRF1 (EA-D) was used as a marker for replication compartments. Cells transfected with SV40p-Cntl (SVpE) or pLVX-Cntl served as negative controls.

We next asked whether BRRF1-induced splicing changes depend on components of the CCR4-NOT complex. DG75 cells were first transfected with either a control siRNA or an siRNA targeting the scaffold subunit CNOT1, and 48 hours later, these cells were co-transfected with either a control or BRRF1 expression vector together with a constitutive GFP expression vector (**Supplemental Figure S6A**, *left panel*). GFP-positive cells were isolated by FACS, and RNA was prepared from sorted cells. RT-qPCR confirmed a 66-81% reduction in CNOT1 expression under knockdown conditions (**Supplemental Figure S6B**). Poly(A)+ RNA sequencing was then performed, and BRRF1-induced splicing changes were compared in the presence or absence of CNOT1 knockdown. Depletion of CNOT1 led to a significant reduction in the number of BRRF1-mediated increased SE events (**Figure 6B**, *left panel*), as well as a significant decrease in the average amplitude of these exon-skipping changes (**Figure 6B**, *right panel*). These findings indicate that the CCR4-NOT complex, via CNOT1, contributes to BRRF1-induced exon skipping.

Because CCR4-NOT is a multi-subunit complex, we next asked whether depletion of additional subunits similarly alters BRRF1-induced splicing changes. We therefore knocked down the deadenylase subunit CNOT6 and the regulatory subunit CNOT9 in DG75 cells expressing BRRF1. RT-qPCR confirmed a 71-76% reduction in CNOT6 expression and a 53-55% reduction in CNOT9 expression relative to controls (**Supplemental Figure S6C and S6D**). Whereas depletion of CNOT6 had little effect on the number and amplitude of significant increased SE events, depletion of CNOT9 resulted in a reduction in both the number of significant increased SE events and the amplitude of increased exon-skipping changes (**Figure 6C and 6D**). Together, these data indicate that BRRF1-induced splicing disruption depends selectively on components of the CCR4-NOT complex, with CNOT1 and CNOT9 playing particularly important roles.

### A conserved CIY(Y/E) motif in BRRF1 mediates CNOT9 binding and BRRF1-induced splicing disruption

Having implicated the CCR4-NOT complex in BRRF1-mediated splicing disruption, we next sought to define the underlying interaction interface. Using AlphaFold3 to predict interactions between BRRF1 and major CCR4-NOT components, we found that CNOT9 showed the highest-confidence predicted interaction with BRRF1 (**Supplemental Table S2**). Visualization of the predicted BRRF1-CNOT9 interface indicated that BRRF1 tyrosine 224 is positioned within one of the hydrophobic clefts of CNOT9 known to engage GW182 family proteins, with possible additional contribution from tyrosine 225 (**Figure 8A**). Notably, despite only 26.3% amino acid identity between BRRF1 and its KSHV counterpart ORF49, tyrosine 224 lies within a conserved CIY(Y/E) motif (BRRF1, CIYY; ORF49, CIYE) (**Supplemental Figure S7A and S7B**). AlphaFold3 similarly predicted a strong interaction between the corresponding ORF49 motif and CNOT9 (**Supplemental Table S2**). These findings suggest that BRRF1 engages the CCR4-NOT complex, at least in part, through interaction of its conserved CIY(Y/E) motif with CNOT9.

**Figure 8.**
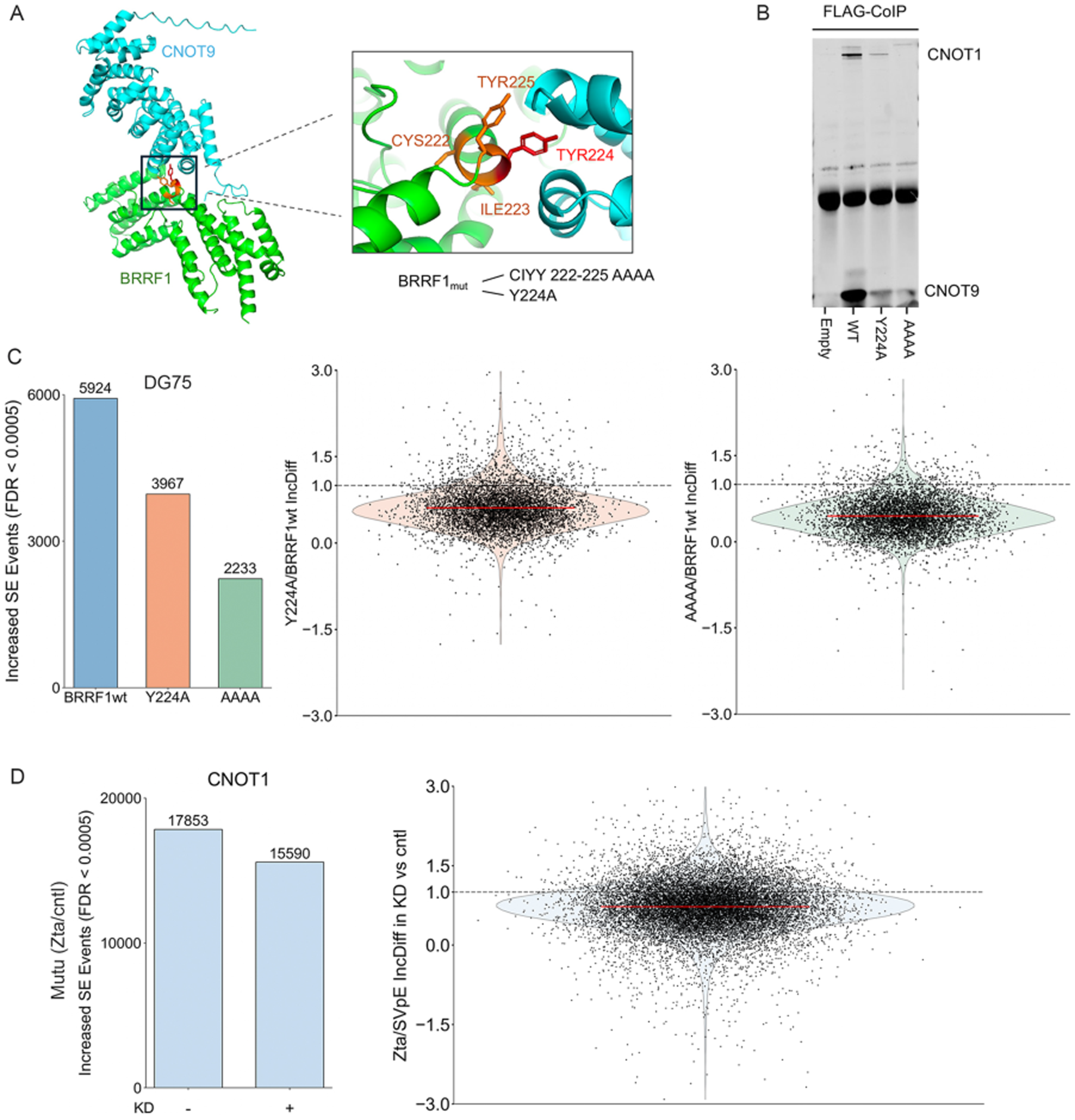
BRRF1’s interaction with the CCR4-NOT complex causes splicing disruption. **(A)** The BRRF1-CNOT9 interacting region was modeled using AlphaFold3, which identified an interacting CIYY motif at positions 222-225, with Y224 predicted to be a critical residue interacting with CNOT9. BRRF1 mutants, BRRF1_Y224A_ (Y224A) and BRRF1_C222A-I223A-Y224A-Y225A_ (AAAA) were generated by substituting the codons for the indicated amino acid positions with alanine codons. **(B)** Validation of the interaction between BRRF1 mutants and CNOT9 and CNOT1 by co-immunoprecipitation. **(C)** SE changes caused by wildtype (WT) and mutant BRRF1. The number of statistically significant increased SE events induced by BRRF1-WT and BRRF1 mutants (bar plots), and the amplitude of increased SE changes following mutation (violin plots), were assessed using rMATS (IncDiff_threshold_ = 0 and FDR < 0.0005). For amplitude analysis, IncDiff values for all significantly increased SE events identified under BRRF1-WT (BRRF1-WT vs. pLVX) were extracted. Corresponding IncDiff values for the same SE events were obtained from each BRRF1 mutant (Y224A or AAAA vs. pLVX). Violin plots show the ratio of IncDiff values for each mutant relative to BRRF1-WT. (**D**) CNOT1 contributes to splicing changes during reactivation. The number of statistically significant increased SE events in the mock (siCntl-Zta vs. siCntl-SVpE) and CNOT1 knockdown (siCNOT1-Zta vs. siCNOT1-SVpE) conditions during Zta-induced reactivation (IncDiff_threshold_ = 0 and FDR < 0.0005) (see **Supplemental Figure S6A,** *right panel*) is shown as bar plot. The amplitude of reactivation-induced SE changes following CNOT1 knockdown was assessed as described in **Figure 6B**, 6D and **8C** and is shown as violin plots.

To test this model experimentally, we generated BRRF1 mutant expression vectors in which either tyrosine 224 was replaced by alanine (BRRF1_Y224A_) or the entire CIYY motif was mutated (BRRF1_C222A-I223A-Y224A-Y225A_) (**Figure 8A**). These substitutions were not predicted to alter the overall BRRF1 fold, as AlphaFold3 predicted both mutant structures with high confidence (pTMs = 0.8, **Supplemental Table S2**), and alignment with wild-type BRRF1 yielded low root mean square deviation (RMSD) values (<0.5 Å) (**Supplemental Figure S7C**). In contrast, AlphaFold3 interaction scores for binding of wild-type BRRF1, BRRF1_Y224A_, and BRRF1_C222A-I223A-Y224A-Y225A_ to CNOT9 were 0.75, 0.12, and 0.27, respectively, suggesting that both mutations weaken binding to CNOT9 (**Supplemental Table S2**).

We next transfected wild-type and mutant BRRF1 expression vectors into DG75 cells, immunoprecipitated BRRF1 with an anti-FLAG antibody, and assessed co-precipitation of CNOT9 and CNOT1 by Western blot. As shown in **Figure 8B**, mutation of Y224 alone significantly reduced the association of BRRF1 with both CNOT9 and CNOT1, whereas the BRRF1_C222A-I223A-Y224A-Y225A_ mutant displayed a near-complete loss of binding (**Figure 8B**). The loss of interaction was not attributable to reduced BRRF1 stability, as both mutants were expressed robustly at levels comparable to wild-type BRRF1 (**Supplemental Figure S7D**). Together, these results experimentally validate the predicted BRRF1-CNOT9 interaction interface and provide reagents to test the role of this interaction in BRRF1-mediated splicing disruption.

To determine whether BRRF1-mediated splicing disruption depends on its interaction with CNOT9, wild-type and mutant BRRF1 expression vectors were co-transfected with a GFP expression vector into DG75 cells, and GFP-positive cells were isolated by FACS. RNA sequencing analysis of sorted cells showed that both mutations reduced the number of increased SE events, with mutation of the entire CIYY motif decreasing BRRF1-mediated exon skipping by 62.3% (**Figure 8C**). In addition, the amplitude of increased exon-skipping events induced by these mutants was significantly reduced (**Figure 8C**). These findings indicate that the CIYY motif is critical for BRRF1 interaction with CNOT9, and that this interaction makes a substantial contribution to BRRF1-mediated splicing disruption.

### CNOT1 contributes to EBV reactivation-associated splicing disruption

Given the role of the CCR4-NOT complex in BRRF1-induced splicing disruption, we asked whether this complex contributes to splicing disruption during EBV reactivation. To address this, we assessed splicing changes in Mutu cells carrying the pCEP4-BMRF1p-GFP lytic reporter following CNOT1 knockdown. Cells were transfected with either a control or CNOT1-specific siRNA 24 hours before induction of reactivation by transfection of a Zta expression vector (**Supplemental Figure S6A**, *right panel*). RT-qPCR confirmed an 80-90% reduction in CNOT1 expression under knockdown conditions (**Supplemental Figure S6E**). Assessment of reactivation-associated splicing changes in this model showed that CNOT1 knockdown caused only a modest reduction in the total number of statistically significant increased SE events, but it substantially reduced the magnitude of increased exon skipping across events (**Figure 8D**). Thus, although many exon-skipping events remained statistically significant after CNOT1 depletion, their severity was broadly attenuated. The persistence of many statistically significant events after CNOT1 knockdown likely reflects the strength of the underlying reactivation-associated splicing program together with incomplete suppression of CCR4-NOT function. These findings indicate that CNOT1 knockdown does not abolish reactivation-associated exon skipping, but significantly dampens its extent, supporting an important contributory role for the CCR4-NOT complex in splicing disruption during EBV reactivation.

## Discussion

### Splicing disruption is a conserved and pervasive feature of gammaherpesvirus reactivation

Herpesviruses have evolved conserved mechanisms to redirect host cell machinery toward productive viral replication while limiting antiviral defenses. Our findings suggest that widespread disruption of host splicing should now be considered part of this conserved host-remodeling repertoire. Using a rigorously controlled and deeply sequenced set of EBV and KSHV reactivation models spanning both lymphoid and epithelial cell contexts, we found that lytic replication is associated with extensive splicing disruption, particularly increased exon skipping, across all systems examined. The substantial overlap in exon-skipping events across models further argues that this is not a model-specific consequence of reactivation, but a conserved feature of gammaherpesvirus lytic replication. A recent report from the Johannsen laboratory describing increased exon skipping and intron retention during EBV reactivation in a lymphoblastoid cell line provides additional support for this conclusion (2).

### The splicing phenotype during reactivation is highly penetrant and predominantly disruptive

A central implication of our findings is that the splicing changes induced during EBV reactivation are more consistent with a broad disruption of splicing fidelity than with selective reprogramming of physiological splicing pathways. Several features of the data support this interpretation. First, exon skipping was the dominant splicing abnormality across all EBV and KSHV reactivation models and reached a magnitude exceeding that observed following knockdown of any of 186 RNA-binding proteins and splicing factors in our comparative dataset. Second, the penetrance of this phenotype was striking, with increased exon skipping affecting up to 56.6% of expressed genes in some reactivation models, indicating that the effects of lytic replication extend across a remarkably large fraction of the host transcriptome. Third, a high proportion of induced exon-skipping events were not represented in existing transcript annotations, arguing against simple engagement of established physiological splicing programs and instead supporting the generation of aberrant splice isoforms during reactivation. Finally, many of these induced exon-skipping events were predicted either to trigger nonsense-mediated decay or to remove coding sequences corresponding to conserved protein domains, suggesting that the net effect of this process is often loss of productive host gene output rather than diversification of functional isoforms. Together, these observations argue that EBV reactivation imposes a widespread loss of host splicing fidelity that is likely to compromise productive cellular gene expression on a transcriptome-wide scale.

### The scale of splicing disruption during reactivation creates broad vulnerability across host pathways

An important implication of the scale of EBV-induced splicing disruption is that its biological effects need not arise from selective targeting of particular cellular pathways. Rather, the breadth and penetrance of the exon-skipping phenotype suggest that EBV may simply impose a widespread loss of splicing fidelity across the host transcriptome, with the consequence that numerous cellular pathways become vulnerable to functional compromise. In this model, pathway alterations that are especially important for the lytic program, including those governing cell cycle progression, innate immune signaling, and RNA processing itself, would be among those affected simply because of the extensive scope of the disruption.

Consistent with this view, several of the exon-skipping events identified here are predicted to compromise pathways that would be expected to influence the efficiency of the lytic cycle. Exon-skipping events affecting CDK2 and CCNE1, for example, are predicted to weaken interactions required for passage through the G1/S transition and progression through S phase, raising the possibility that splicing disruption contributes to reinforcement of the G1/S-like state previously associated with EBV lytic replication. Innate immune signaling pathways may likewise be especially vulnerable to this type of transcriptome-wide disruption. In this regard, it is notable that many interferon genes are themselves intronless, which may represent an evolutionary strategy to preserve rapid antiviral gene induction under conditions in which splicing is impaired. By contrast, many upstream regulators and downstream interferon response genes are multi-exonic and therefore potentially susceptible to the widespread exon skipping observed here. Although our analyses could not directly detect aberrant splicing of inducible interferon-stimulated transcripts that are not expressed before pathway activation, the highly penetrant nature of the splicing disruption suggests that many such transcripts would also be vulnerable to production of functionally compromised isoforms once induced. Nevertheless, sensors and regulators of innate immune signaling are broadly impacted by splicing disruption including reactivation-associated exon skipping in TYK2 that is predicted to remove critical kinase-associated domains and thereby impair antiviral signaling through a dominant negative action. Finally, the enrichment of exon-skipping events in RNA metabolism pathways, including factors involved in RNA processing and splicing, raises the possibility of a self-reinforcing cascade in which early disruption of host splicing further compromises the machinery needed to maintain normal RNA processing. Together, these observations suggest that large-scale splicing disruption may help create a cellular environment that is permissive for viral replication while simultaneously weakening antiviral defenses.

### Early splicing disruption is poised to preempt host signaling responses

The minimal effect of phosphonoacetic acid on the number, character, and amplitude of reactivation-associated splicing changes argues that this phenotype arises largely independently of viral DNA replication and late gene expression. Instead, the data place the onset of splicing disruption early in the lytic cascade, indicating that it is part of the host-remodeling program rather than a secondary consequence of late lytic events. This timing is likely advantageous to EBV, as early disruption of host splicing would impair the cell’s ability to mount new transcriptional responses before those programs are fully engaged. Such an effect may be especially important for pathways that must be induced rapidly following reactivation, including innate immune, stress-response, and cell cycle regulatory programs. Early splicing disruption may likewise reinforce the G1/S-like cellular state associated with EBV lytic replication and, through effects on transcripts encoding RNA-processing factors, initiate a self-reinforcing decline in host RNA-processing capacity. More broadly, to the extent that the viral lytic transcriptome is less dependent on splicing than the host transcriptome, this strategy would be expected to disproportionately impair host gene expression while leaving viral gene expression relatively less affected. Together, these considerations argue that early splicing disruption is likely to be advantageous because it destabilizes host gene-expression programs before later stages of the lytic cycle dominate.

### BRRF1 recapitulates key features of reactivation-associated splicing disruption

If splicing disruption is established early during reactivation, then one or more early viral gene products are likely to initiate or amplify this phenotype. Our early-gene screen showed that several EBV early genes can influence splicing, but BRRF1 stood out as the factor whose phenotype most closely resembled the disruptive pattern observed during reactivation. BRRF1 was distinguished by its strong bias toward increased exon skipping, its high penetrance across expressed genes, its enrichment for unannotated exon-skipping events, and its ability to induce intron retention alongside exon skipping. Most importantly, BRRF1 showed the strongest overlap with the reactivation-associated exon-skipping program across the EBV models tested, arguing that it is not simply altering splicing in a general way, but instead recapitulating a substantial component of the specific disruptive phenotype seen during lytic induction. These findings support the view that BRRF1 functions less as a pathway-specific splicing regulator and more as an early amplifier of host permissiveness by promoting broad loss of splicing fidelity. At the same time, our data do not suggest that BRRF1 is the sole determinant of this phenotype, but rather an important contributor within a likely multifactorial early lytic program. This interpretation fits well with the broader attenuation-without-complete-abrogation pattern seen elsewhere in the study, which argues for contribution by multiple viral and host factors rather than control by a single dominant effector. Thus, BRRF1 emerges as the strongest early-gene candidate linking early viral gene expression to the splicing-disruption phenotype observed during EBV reactivation.

This interpretation is also consistent with prior work on BRRF1, which has suggested that its contribution to the lytic cycle is context dependent rather than universally required. In some epithelial systems, BRRF1 enhances BRLF1-mediated lytic induction, and can also independently induce the expression of several lytic genes (66,67). On the contrary, BRRF1-deficient virus shows no obvious defect in viral replication in HEK293 cells (68). Our findings suggest one way to reconcile these observations. The advantage conferred by BRRF1-mediated splicing disruption may be most apparent in settings where rapid suppression of inducible host signaling pathways, including innate immune responses, is important, a pressure that is likely to be muted in standard in vitro systems. In addition, the host pathways affected by splicing disruption are unlikely to depend on this mechanism alone. For example, disruption of transcripts such as CCNE1 and CDK2 may reinforce a G1/S-like state that is also promoted through other viral activities, such that loss of BRRF1 would diminish this layer of support without abolishing the overall cellular state needed for productive replication. In this view, BRRF1-dependent splicing disruption would function less as an essential switch and more as an early reinforcing mechanism that helps stabilize a host environment favorable for the lytic program.

### EBV exploits an underappreciated nuclear function of the CCR4-NOT complex

Our findings identify the CCR4-NOT complex as a likely mechanistic bridge between BRRF1 expression and reactivation-associated splicing disruption. This is notable because CCR4-NOT is best known as a cytoplasmic regulator of mRNA fate, particularly in deadenylation and turnover (65,69,70), rather than as a canonical splicing factor. However, the complex has also been implicated in nuclear gene-regulatory functions, and some of its subunits appear to serve important recruitment roles in addition to supporting catalytic activity. In particular, CNOT9 is a known interaction surface through which cellular factors such as GW182 and tristetraprolin (TTP) recruit or engage the CCR4-NOT complex (71–74), making it plausible that BRRF1 could use this same interface to redirect the complex toward a distinct function. Consistent with this idea, our data show that BRRF1 associates with CNOT1 and CNOT9 in the nucleus (**Figure 7A and 7B**), and that BRRF1 is predicted to engage CNOT9 through the same hydrophobic cleft used by GW182-family proteins (71,72). Together, these findings suggest that EBV may exploit and/or recruit a nuclear, recruitment-competent pool of the CCR4-NOT complex to compromise host splicing fidelity during reactivation.

The subunit-selective nature of our results has mechanistic implications. Depletion of CNOT1 and CNOT9 attenuated BRRF1-induced exon skipping, whereas depletion of the deadenylase subunit CNOT6 had little effect. This pattern argues against a model in which BRRF1 disrupts splicing simply by engaging generic CCR4-NOT deadenylase activity (although we have not knocked down CNOT6 and CNOT6L and/or CNOT7/8 simultaneously). Instead, it supports a mechanism in which BRRF1 engages the recruitment and organizational functions of the complex, centered on CNOT1 and CNOT9. That interpretation is reinforced by the CIYY-motif data: BRRF1 uses a conserved CNOT9-contacting interface, mutation of that interface weakens BRRF1 association with both CNOT9 and CNOT1, and those mutations substantially reduce BRRF1-mediated exon skipping. These findings raise the possibility that BRRF1 hijacks a normal CCR4-NOT recruitment surface and retargets the complex toward a nuclear activity that destabilizes host splicing fidelity. In this view, CNOT9 may provide the molecular entry point for BRRF1 engagement, while CNOT1 supplies the broader complex architecture needed to confer that interaction into functional effects on host RNA processing.

Our data do not yet define exactly how BRRF1-bound CCR4-NOT disrupts splicing, but they do narrow the plausible models. One possibility is that BRRF1 engagement of nuclear CCR4-NOT alters transcriptional context, such as RNA polymerase II elongation dynamics or other cotranscriptional RNA-processing events, thereby secondarily biasing splice-site choice toward exon skipping and intron retention. This model is attractive because CCR4-NOT has recognized links to transcription initiation and elongation (63,65,75,76), and transcriptional dynamics are well positioned to influence splice-site recognition on a transcriptome-wide scale. A second, nonexclusive possibility is that BRRF1-bound CCR4-NOT perturbs the assembly, recruitment, or stability of RNA-processing complexes required for normal exon definition, particularly at splice-acceptors that are already vulnerable to disruption. Either mechanism would be consistent with the unusually broad, highly penetrant, and predominantly disruptive splicing phenotype observed during reactivation.

CCR4-NOT may be an especially effective host target for EBV because it influences multiple layers of gene-expression control rather than a single isolated process. In addition to its established role in mRNA decay, the complex has been linked to transcriptional regulation and other aspects of RNA metabolism, making it a multifunctional control point in host transcriptome regulation. By perturbing such a hub, EBV could generate broad downstream vulnerability across the host transcriptome without having to target many cellular pathways individually. This interpretation fits well with the broader picture that emerges from our study: rather than using splicing to selectively rewire a small number of pathways, EBV appears to impose a high-penetrance loss of splicing fidelity with widespread collateral effects on host gene expression. This also raises the possibility that BRRF1 engagement of CCR4-NOT extends beyond splicing disruption to other functions of the complex in host transcriptome regulation.

The relevance of the BRRF1-CCR4-NOT axis to EBV reactivation is strengthened by our finding that CNOT1 knockdown in reactivating Mutu cells attenuated the amplitude of exon skipping, even though many events remained statistically significant. That pattern is consistent with CCR4-NOT acting as an important organizer or amplifier of the phenotype in the native reactivation setting. More broadly, these findings suggest that BRRF1 engagement of CCR4-NOT represents a major mechanistic axis of the splicing-disruption program during reactivation, even if additional viral or host factors help shape its full extent. Finally, the conservation of the CNOT9-contacting CIY(Y/E) motif between BRRF1 and KSHV ORF49 raises the possibility that engagement of CCR4-NOT-related functions may reflect a more general gammaherpesvirus strategy. Defining how BRRF1 retargets CCR4-NOT should therefore help clarify whether this mechanism operates primarily through transcription-coupled RNA processing, altered assembly of RNA-processing machinery, or a combination of both.

## Data availability

RNA sequencing datasets for this study are available at the NCBI Geo Archive with the accession number, GSE240008.

## Acknowledgements

We acknowledge the support of the computational resources and expertise of the Tulane Cancer Center Next Generation Sequence Analysis Core. The content of this article is solely the responsibility of the authors and does not necessarily represent the views of the funding agencies.

## Author contributions

T.T.N., T.D.N., and E.K.F. designed the study; T.T.N., T.D.N., A.G., T.M.O., C.R., E.I., N.W., J.B., M.L., M.B. and N.V.O. performed experiments; Q.Z., H.L., N.A.U., R.R., and Y.D. provided consultations on experimental approaches and design; T.T.N., T.D.N., and E.K.F. wrote the manuscript.

## Declaration of generative AI and AI-assisted technologies in the writing process

The authors utilized ChatGPT 5.4 to enhance grammar and readability during the manuscript writing. After this, the authors thoroughly reviewed and edited the article, assuming full responsibility for its content.

## Supplemental Figure Legends

**Supplemental Figure S1.**
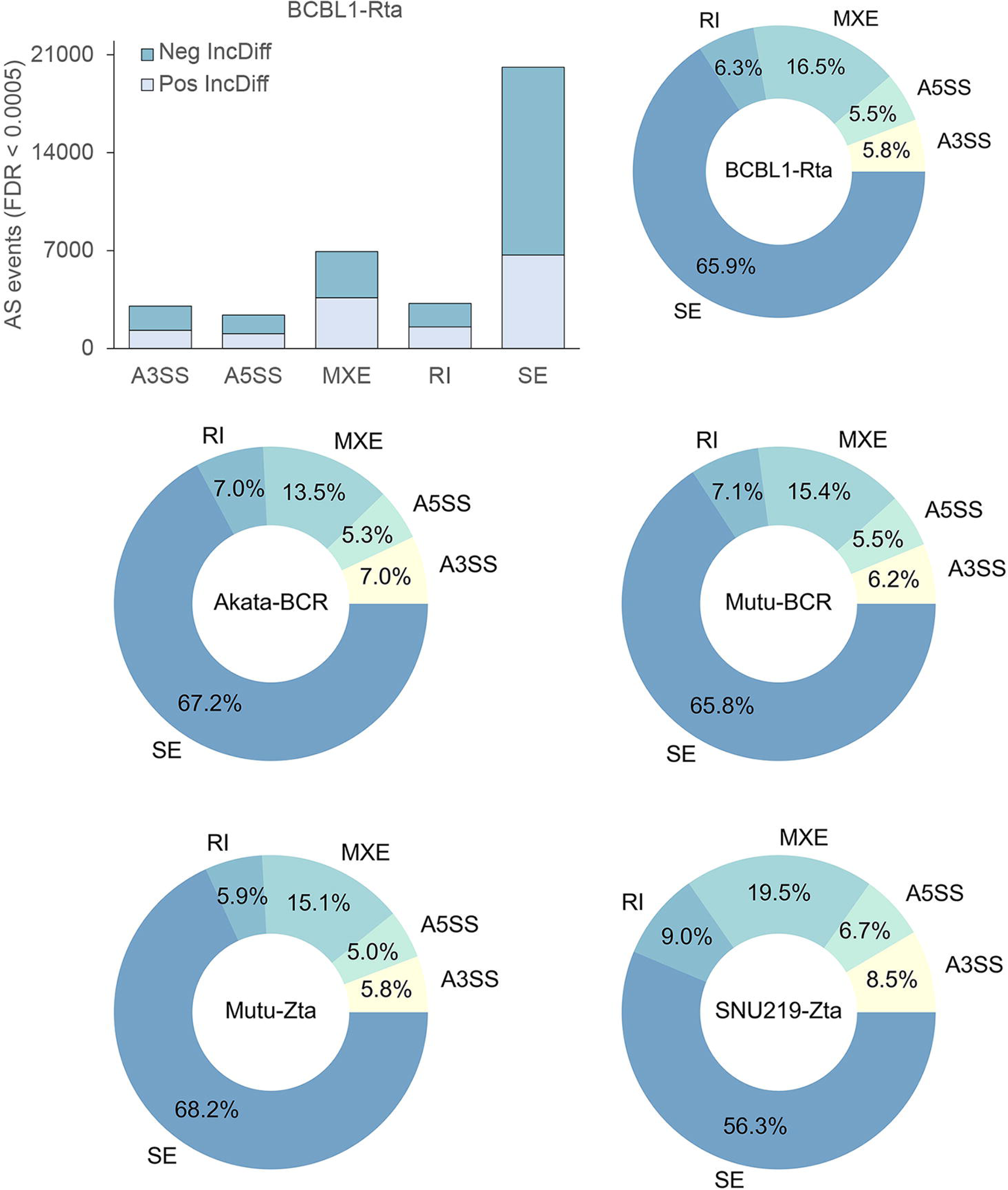
Differential alternative splicing analyses of EBV and KSHV reactivations. The number of differential AS events in KSHV reactivation model (FDR < 0.0005) is shown as a bar plot. The percentage of statistically significant events (Neg-IncDiff and Pos-IncDiff) for each AS type relative to the total number of significant events across all five AS types is shown as pie charts for EBV and KSHV reactivation models (FDR < 0.0005).

**Supplemental Figure S2.**
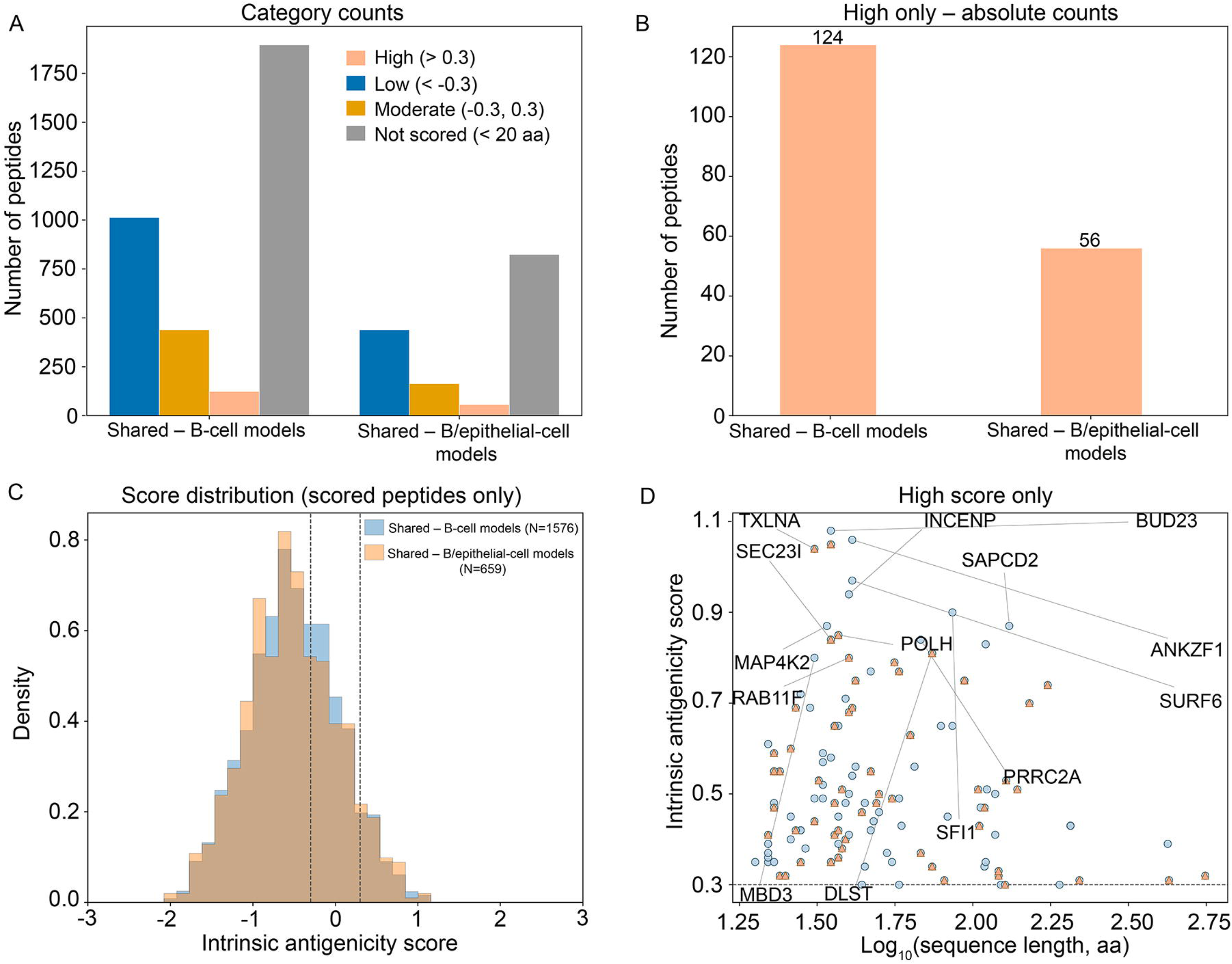
Antigenicity analysis of reactivation-derived neopeptides. The neopeptides shared among our EBV+ B cell, and B and epithelial cell reactivation models were analyzed for their predicted antigenicity using IAPred (38). (**A**) The number of peptides that were predicted to have high (> 0.3), low (< −0.3), or moderate (−0.3 to 0.3) for B cell and B and epithelial cell reactivation models. (**B)** The number of neopeptides predicted to have high antigenicity shared among B and among B and epithelial cell reactivation models. (**C**) Density distribution of intrinsic antigenicity scores for neopeptides shared among B cell reactivation models (blue) and shared B and epithelial cell reactivation models (orange). (**D**) Correlation between intrinsic antigenicity score and neopeptide length for high-scoring neopeptides (> 0.3). Each point represents a neopeptide shared among B cell reactivation models (blue) and among B and epithelial cell reactivation models (orange). Top 15 selected genes corresponding to the highest antigenicity scores are indicated.

**Supplemental Figure S3.**
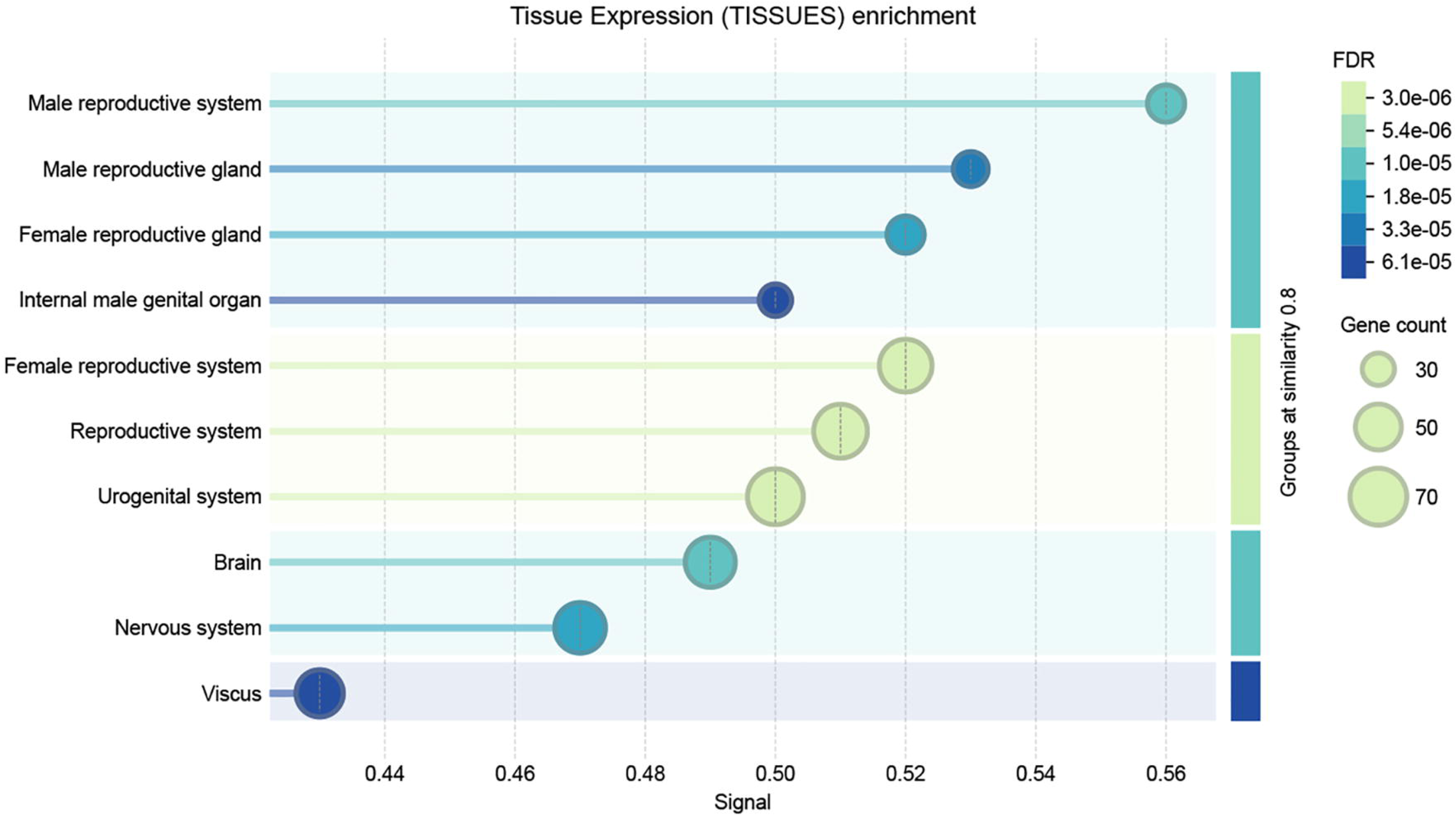
Tissue enrichment analysis of genes corresponding to highly antigenic neopeptides. Genes corresponding to predicted highly antigenic neopeptides were analyzed for tissue enrichment using STRING (39). Enriched gene sets were considered statistically significant at FDR < 0.05.

**Supplemental Figure S4.**
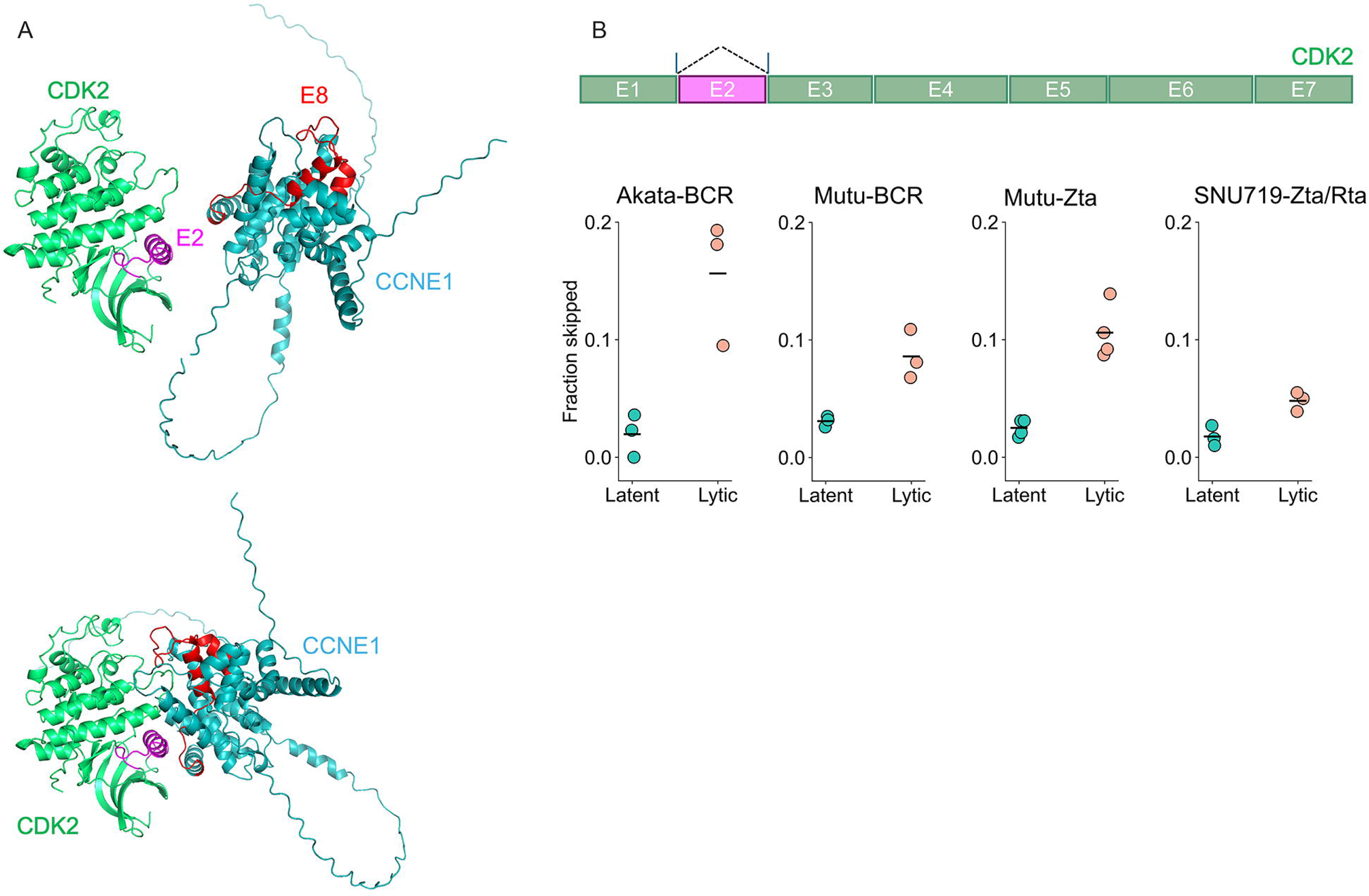
Significant exon 2 skipping in CDK2. **(A)** A 3D structural model of the CDK2-CCNE1 complex was predicted using AlphaFold3 and visualized in PyMOL. Exon 2 in CDK2 (magenta) and exon 8 in CCNE1 (red) are indicated. In addition to the skipped exon regions, other regions in both CDK2 and CCNE1 appear to contribute to the interaction interface. This model was used to identify interface residues involved in the CDK2-CCNE1 interaction. **(B)** The fraction of exon 2 skipping in CDK2 was calculated using the same approach as for CCNE1 and TYK2 in **Figure 4**.

**Supplemental Figure S5.**
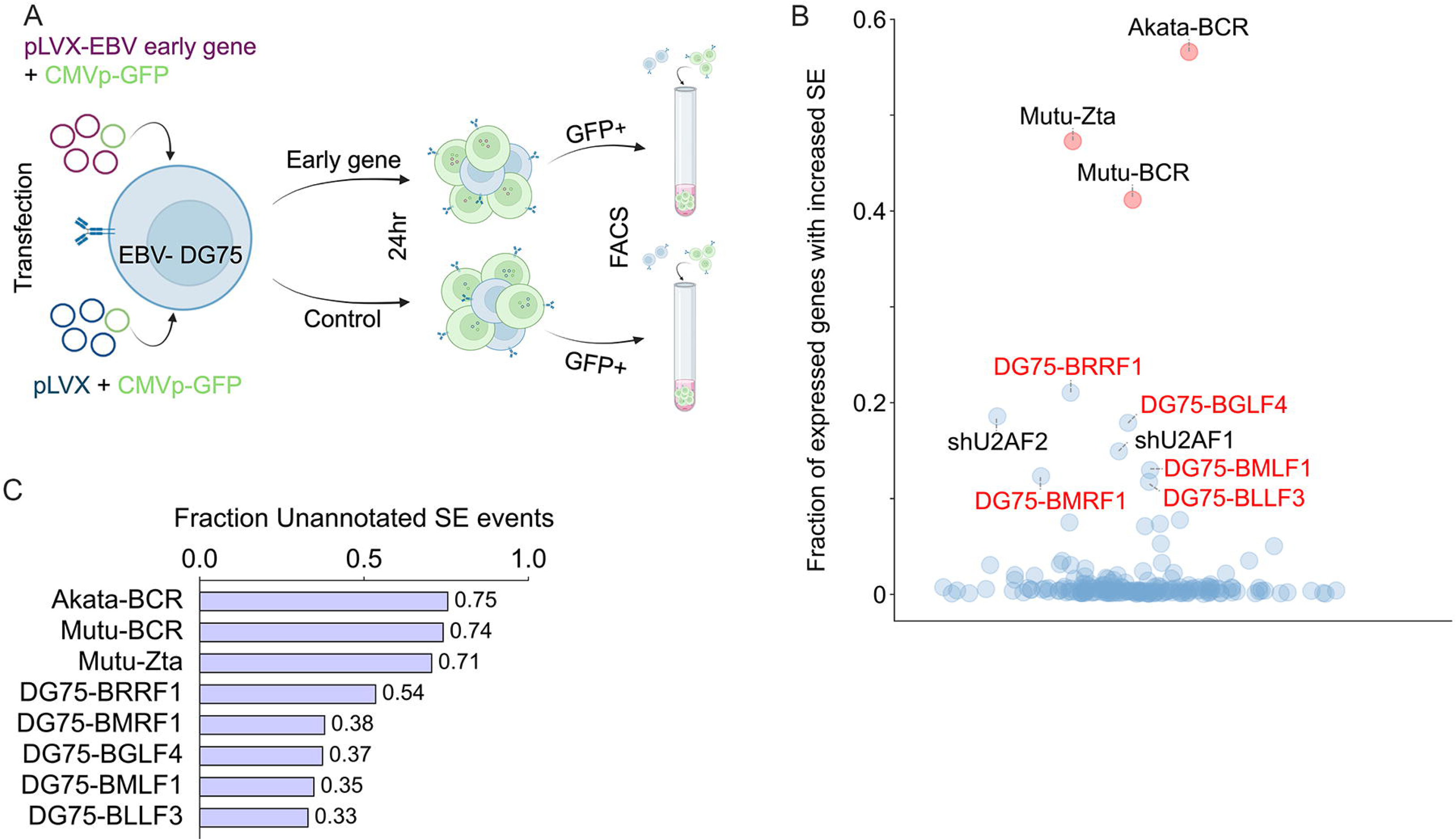
BRRF1 exhibits splicing-disrupting phenotype. (**A**) Workflow of systemic analysis of the effect of each EBV early gene expression on cell splicing changes. The EBV-negative Burkitt lymphoma cell line DG75 was co-transfected with CMVp-GFP and a pLVX plasmid expressing either a control or each EBV early gene open reading frame (ORF). After 24 hours, GFP+ cells were selected from each sample by FACS (N=3 or 4 per group). The panel was created using BioRender.com.sarala(**B**) Fraction of expressed genes exhibiting increased exon skipping in EBV reactivation models, the top five EBV early genes with the most significant influence on splicing changes, and knockdowns of 186 RNA-binding proteins (RBPs) using SEFractionExpressed module (SpliceTools; FDR < 0.0005; TPM ≥ 3 in either condition).sarala(**C**) Fraction of increased SE events that are unannotated in EBV reactivation models and in the top five EBV early genes with the greatest impact on increased exon skipping, quantified using SEUnannotated module (SpliceTools; FDR < 0.0005).

**Supplemental Figure S6.**
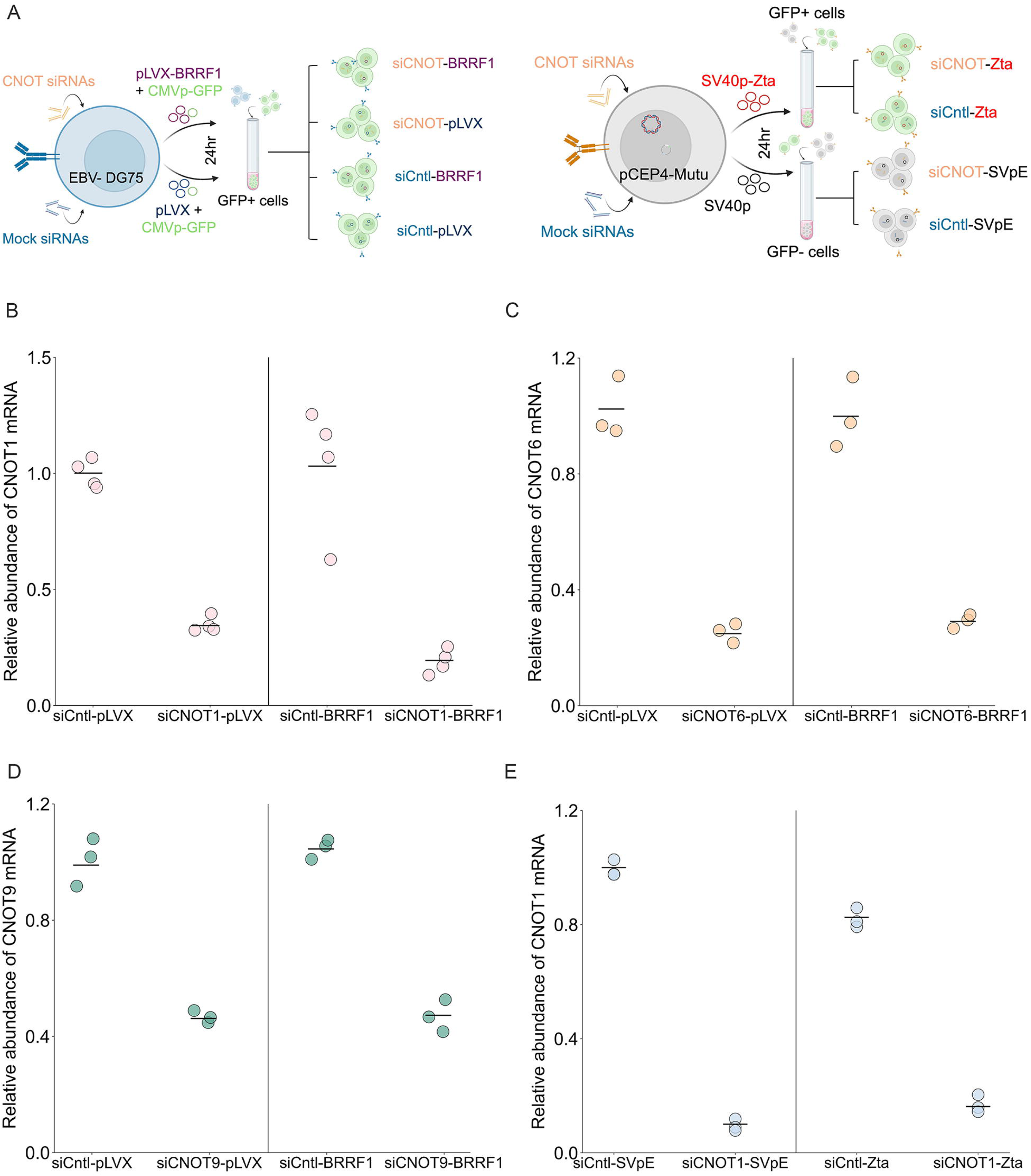
Knockdown of CCR4-NOT subunits. (**A**) Workflow of CCR4-NOT subunit knockdowns in DG75 (*left panel*) and pCEP4-BMRF1p-GFP Mutu cells (*right panel*). (*left panel*) DG75 cells were transfected with either mock siRNAs (siCntl) or CNOT-specific siRNAs (siCNOT) for 48 hours (CNOT1), 6 hours (CNOT6), or 18 hours (CNOT9). The samples were then split into two groups, one that was co-transfected with the CMVp-GFP plasmid along with either the BRRF1 expression plasmid or the control (pLVX) plasmid. This produced four experimental conditions: siCntl-pLVX, siCntl-BRRF1, siCNOT-pLVX, and siCNOT-BRRF1. After 24 hours, GFP+ cells were sorted using FACS. (*right panel*) A similar experimental setup was conducted in pCEP4-BMRF1p-GFP Mutu cells. After 24 hours of the first transfection with either mock or CNOT1-specific siRNAs, the samples were split into two groups. One group was reactivated with SV40p-Zta, while the other group was transfected with SV40p-Cntl (SVpE) for 24 hours. This setup resulted in four experimental conditions as shown in the figure. GFP+ and GFP-cells were sorted using FACS. The panel was created with Biorender.com. (**B-E**) RT-qPCR analysis for knockdown of CCR4-NOT subunits. CNOT1 (**B**), CNOT6 (**C**), and CNOT9 (**D**) knockdown in DG75 and CNOT1 knockdown in pCEP4-BMRF1p-GFP Mutu (**E**).

**Supplemental Figure S7.**
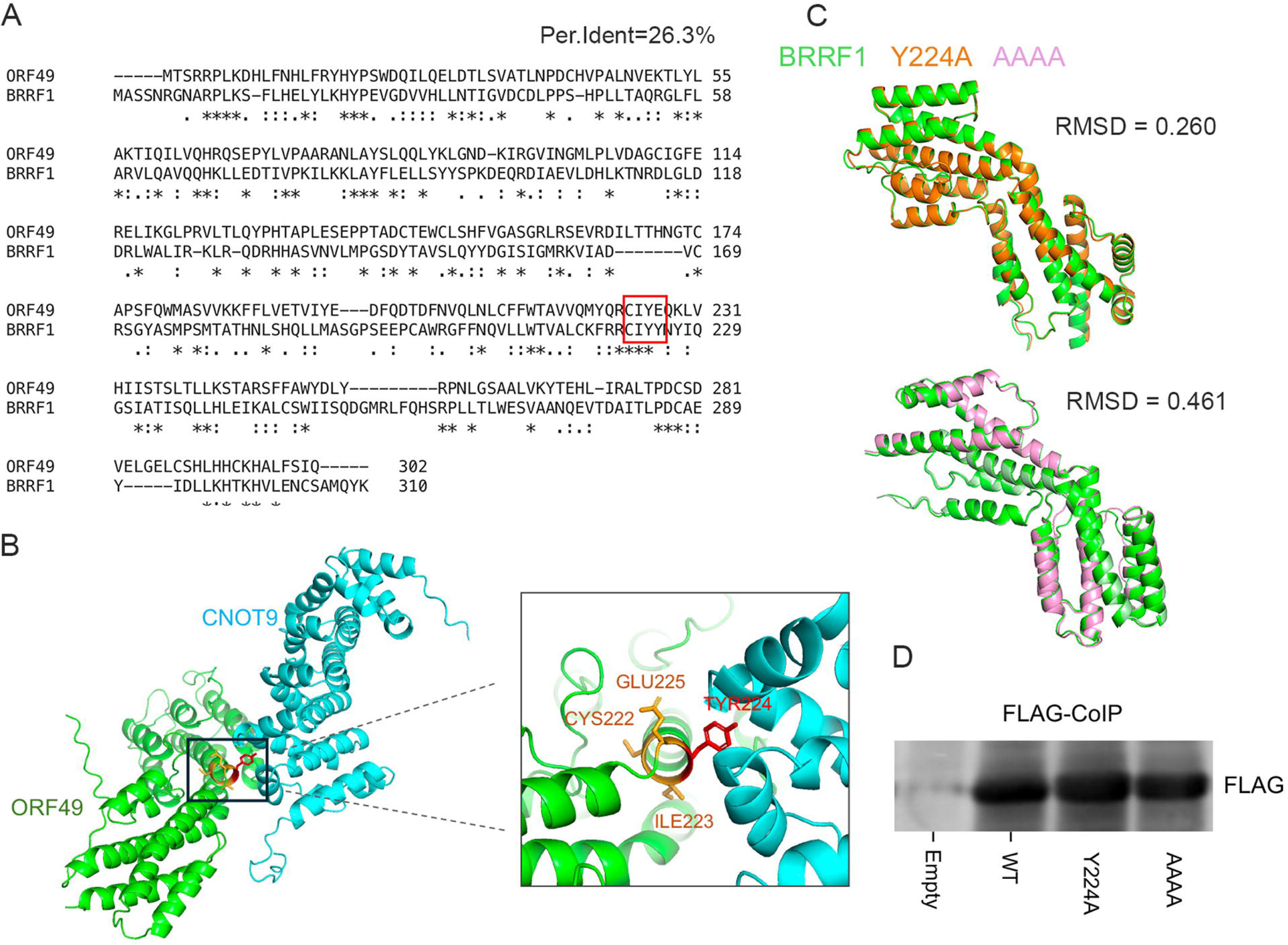
Sequence and structural homology between BRRF1 and KSHV ORF49. (**A**) Protein sequences of EBV BRRF1 and its homolog in KSHV, ORF49, were aligned using Clustal Omega (36). Identical residues are indicated by (*). The percentage sequence identity between BRRF1 and ORF49 was calculated using BLASTp suite (NCBI) (37). (**B**) The ORF49-CNOT9 interaction interface was modeled using AlphaFold3. The CIYE motif (Cysteine-Isoleucine-Tyrosine-Glutamic acid) at the position 224-227, which is highly conserved with the CIYY motif in BRRF1, is predicted to contribute to CNOT9 binding. Similarly to BRRF1, tyrosine 226 in ORF49 is predicted to be a critical residue mediating interaction with CNOT9. (**C**) The expression level of BRRF1 mutants. From the co-IP experiment performed to validate the interaction between BRRF1 mutants and CNOT9, anti-FLAG antibody was used to assess the expression levels of BRRF1 mutants relative to BRRF1-WT. (**D**) Structural homology analysis of BRRF1-WT and mutants. Structure of BRRF1-WT and BRRF1 mutants was predicted individually using Alphafold3. Each mutant structure was aligned to BRRF-WT in PyMOL, which gave root-mean-square deviation (RMSD) values to assess structural similarity. An RMSD < 2Å indicates high structural similarity.

## Reference

1. Van Nostrand, E.L., Freese, P., Pratt, G.A., Wang, X., Wei, X., Xiao, R., Blue, S.M., Chen, J.Y., Cody, N.A.L., Dominguez, D., et al. (2020) A large-scale binding and functional map of human RNA-binding proteins. Nature, 583, 711–719.

2. Casco, A., Ohashi, M. and Johannsen, E. (2024) Epstein-Barr virus induces host shutoff extensively via BGLF5-independent mechanisms. Cell Rep, 43, 114743.

3. Ungerleider, N.A., Roberts, C., O’Grady, T.M., Nguyen, T.T., Baddoo, M., Wang, J., Ishaq, E., Concha, M., Lam, M., Bass, J. et al. (2024) Viral reprogramming of host transcription initiation. Nucleic Acids Res, 52, 5016–5032.

4. Rosemarie, Q., Kirschstein, E. and Sugden, B. (2023) How Epstein-Barr Virus Induces the Reorganization of Cellular Chromatin. mBio, 14, e0268622.

5. Verma, D. and Swaminathan, S. (2008) Epstein-Barr virus SM protein functions as an alternative splicing factor. J Virol, 82, 7180–7188.

6. Verma, D., Bais, S., Gaillard, M. and Swaminathan, S. (2010) Epstein-Barr Virus SM protein utilizes cellular splicing factor SRp20 to mediate alternative splicing. J Virol, 84, 11781–11789.

7. Shannon-Lowe, C. and Rickinson, A. (2019) The Global Landscape of EBV-Associated Tumors. Front Oncol, 9, 713.

8. Sugiokto, F.G. and Li, R. (2025) Targeting EBV Episome for Anti-Cancer Therapy: Emerging Strategies and Challenges. Viruses, 17.

9. Meirhaeghe, M.R. and Balfour, H.H., Jr. (2025) Epstein-Barr virus (EBV) infection and its sequelae in the immunocompetent host. J Clin Virol, 180, 105854.

10. Huang, J., Tengvall, K., Lima, I.B., Hedstrom, A.K., Butt, J., Brenner, N., Gyllenberg, A., Stridh, P., Khademi, M., Ernberg, I. et al. (2024) Genetics of immune response to Epstein-Barr virus: prospects for multiple sclerosis pathogenesis. Brain, 147, 3573–3582.

11. Wahbeh, F. and Sabatino, J.J. (2025) Epstein-Barr Virus in Multiple Sclerosis: Past, Present, and Future. Neurol Neuroimmunol Neuroinflamm, 12, e200460.

12. Bjornevik, K., Cortese, M., Healy, B.C., Kuhle, J., Mina, M.J., Leng, Y., Elledge, S.J., Niebuhr, D.W., Scher, A.I., Munger, K.L. et al. (2022) Longitudinal analysis reveals high prevalence of Epstein-Barr virus associated with multiple sclerosis. Science, 375, 296–301.

13. Lyu, L., Li, Q. and Wang, C. (2025) EBV Latency Programs: Molecular and Epigenetic Regulation and Its Role in Disease Pathogenesis. J Med Virol, 97, e70501.

14. Murata, T., Sugimoto, A., Inagaki, T., Yanagi, Y., Watanabe, T., Sato, Y. and Kimura, H. (2021) Molecular Basis of Epstein-Barr Virus Latency Establishment and Lytic Reactivation. Viruses, 13.

15. Damania, B., Kenney, S.C. and Raab-Traub, N. (2022) Epstein-Barr virus: Biology and clinical disease. Cell, 185, 3652–3670.

16. Shannon-Lowe, C., Rickinson, A.B. and Bell, A.I. (2017) Epstein-Barr virus-associated lymphomas. Philos Trans R Soc Lond B Biol Sci, 372.

17. 17. El-Sharkawy, A., Al Zaidan, L. and Malki, A. (2018) Epstein-Barr Virus-Associated Malignancies: Roles of Viral Oncoproteins in Carcinogenesis. Front Oncol, 8, 265.

18. Rosemarie, Q. and Sugden, B. (2020) Epstein-Barr Virus: How Its Lytic Phase Contributes to Oncogenesis. Microorganisms, 8.

19. Dorothea, M., Xie, J., Yiu, S.P.T. and Chiang, A.K.S. (2023) Contribution of Epstein-Barr Virus Lytic Proteins to Cancer Hallmarks and Implications from Other Oncoviruses. Cancers (Basel), 15.

20. Munz, C. (2019) Latency and lytic replication in Epstein-Barr virus-associated oncogenesis. Nat Rev Microbiol, 17, 691–700.

21. Casco, A. and Johannsen, E. (2023) EBV Reactivation from Latency Is a Degrading Experience for the Host. Viruses, 15.

22. Covarrubias, S., Richner, J.M., Clyde, K., Lee, Y.J. and Glaunsinger, B.A. (2009) Host shutoff is a conserved phenotype of gammaherpesvirus infection and is orchestrated exclusively from the cytoplasm. J Virol, 83, 9554–9566.

23. Rowe, M., Glaunsinger, B., van Leeuwen, D., Zuo, J., Sweetman, D., Ganem, D., Middeldorp, J., Wiertz, E.J. and Ressing, M.E. (2007) Host shutoff during productive Epstein-Barr virus infection is mediated by BGLF5 and may contribute to immune evasion. Proc Natl Acad Sci U S A, 104, 3366–3371.

24. Horst, D., Burmeister, W.P., Boer, I.G., van Leeuwen, D., Buisson, M., Gorbalenya, A.E., Wiertz, E.J. and Ressing, M.E. (2012) The “Bridge” in the Epstein-Barr virus alkaline exonuclease protein BGLF5 contributes to shutoff activity during productive infection. J Virol, 86, 9175–9187.

25. Buschle, A., Mrozek-Gorska, P., Cernilogar, F.M., Ettinger, A., Pich, D., Krebs, S., Mocanu, B., Blum, H., Schotta, G., Straub, T. et al. (2021) Epstein-Barr virus inactivates the transcriptome and disrupts the chromatin architecture of its host cell in the first phase of lytic reactivation. Nucleic Acids Res, 49, 3217–3241.

26. Wang, Y., Xie, Z., Kutschera, E., Adams, J.I., Kadash-Edmondson, K.E. and Xing, Y. (2024) rMATS-turbo: an efficient and flexible computational tool for alternative splicing analysis of large-scale RNA-seq data. Nat Protoc, 19, 1083–1104.

27. Flemington, E.K., Flemington, S.A., O’Grady, T.M., Baddoo, M., Nguyen, T., Dong, Y. and Ungerleider, N.A. (2023) SpliceTools, a suite of downstream RNA splicing analysis tools to investigate mechanisms and impact of alternative splicing. Nucleic Acids Res, 51, e42.

28. Chen, E.Y., Tan, C.M., Kou, Y., Duan, Q., Wang, Z., Meirelles, G.V., Clark, N.R. and Ma’ayan, A. (2013) Enrichr: interactive and collaborative HTML5 gene list enrichment analysis tool. BMC Bioinformatics, 14, 128.

29. Kuleshov, M.V., Jones, M.R., Rouillard, A.D., Fernandez, N.F., Duan, Q., Wang, Z., Koplev, S., Jenkins, S.L., Jagodnik, K.M., Lachmann, A. et al. (2016) Enrichr: a comprehensive gene set enrichment analysis web server 2016 update. Nucleic Acids Res, 44, W90–97.

30. Xie, Z., Bailey, A., Kuleshov, M.V., Clarke, D.J.B., Evangelista, J.E., Jenkins, S.L., Lachmann, A., Wojciechowicz, M.L., Kropiwnicki, E., Jagodnik, K.M., et al. (2021) Gene Set Knowledge Discovery with Enrichr. Curr Protoc, 1, e90.

31. Abramson, J., Adler, J., Dunger, J., Evans, R., Green, T., Pritzel, A., Ronneberger, O., Willmore, L., Ballard, A.J., Bambrick, J. et al. (2024) Accurate structure prediction of biomolecular interactions with AlphaFold 3. Nature, 630, 493–500.

32. Schrödinger, LLC. The PyMOL Molecular Graphics System, Version 3.1.6.1.

33. UniProt, C. (2025) UniProt: the Universal Protein Knowledgebase in 2025. Nucleic Acids Res, 53, D609–D617.

34. Blum, M., Andreeva, A., Florentino, L.C., Chuguransky, S.R., Grego, T., Hobbs, E., Pinto, B.L., Orr, A., Paysan-Lafosse, T., Ponamareva, I. et al. (2025) InterPro: the protein sequence classification resource in 2025. Nucleic Acids Res, 53, D444–D456.

35. Mistry, J., Chuguransky, S., Williams, L., Qureshi, M., Salazar, G.A., Sonnhammer, E.L.L., Tosatto, S.C.E., Paladin, L., Raj, S., Richardson, L.J. et al. (2021) Pfam: The protein families database in 2021. Nucleic Acids Res, 49, D412–D419.

36. Madeira, F., Madhusoodanan, N., Lee, J., Eusebi, A., Niewielska, A., Tivey, A.R.N., Lopez, R. and Butcher, S. (2024) The EMBL-EBI Job Dispatcher sequence analysis tools framework in 2024. Nucleic Acids Res, 52, W521–W525.

37. Altschul, S.F., Madden, T.L., Schaffer, A.A., Zhang, J., Zhang, Z., Miller, W. and Lipman, D.J. (1997) Gapped BLAST and PSI-BLAST: a new generation of protein database search programs. Nucleic Acids Res, 25, 3389–3402.

38. Miles, S., Menafra, G., Iriarte, A. & Chabalgoity, J. A. (2025) IApred: A versatile open-source tool for predicting protein antigenicity across diverse pathogens. ImmunoInformatics, 20.

39. Szklarczyk, D., Kirsch, R., Koutrouli, M., Nastou, K., Mehryary, F., Hachilif, R., Gable, A.L., Fang, T., Doncheva, N.T., Pyysalo, S. et al. (2023) The STRING database in 2023: protein-protein association networks and functional enrichment analyses for any sequenced genome of interest. Nucleic Acids Res, 51, D638–D646.

40. Park, A., Oh, S., Jung, K.L., Choi, U.Y., Lee, H.R., Rosenfeld, M.G. and Jung, J.U. (2020) Global epigenomic analysis of KSHV-infected primary effusion lymphoma identifies functional MYC superenhancers and enhancer RNAs. Proc Natl Acad Sci U S A, 117, 21618–21627.

41. Hong, Y., Shekhar, R., McMahon, S., Keil, N., Baddoo, M., Watkins, J.M., Keshishian, H., Stanclift, C., Carr, S.A., Munschauer, M. et al. (2026) Herpesvirus-encoded lncRNA targets host splicing by interacting with splicing factors. PLoS Pathog, 22, e1014072.

42. Borghol, A.H., Bitar, E.R., Hanna, A., Naim, G. and Rahal, E.A. (2025) The role of Epstein-Barr virus in autoimmune and autoinflammatory diseases. Crit Rev Microbiol, 51, 296–316.

43. Gupta, S., Reddy, V., Lodha, L. and Ashwini, M.A. (2026) Epstein-Barr Virus (EBV) and Autoimmune Diseases: Pathogenic Mechanisms and Therapeutic Insights. J Med Virol, 98, e70785.

44. Morawiec, N., Adamczyk, B., Spyra, A., Herba, M., Boczek, S., Korbel, N., Polechonski, P. and Adamczyk-Sowa, M. (2025) The Role of Epstein-Barr Virus in the Pathogenesis of Autoimmune Diseases. Medicina (Kaunas), 61.

45. Yockteng-Melgar, J., Shire, K., Cheng, A.Z., Malik-Soni, N., Harris, R.S. and Frappier, L. (2022) G(1)/S Cell Cycle Induction by Epstein-Barr Virus BORF2 Is Mediated by P53 and APOBEC3B. J Virol, 96, e0066022.

46. Cayrol, C. and Flemington, E.K. (1996) The Epstein-Barr virus bZIP transcription factor Zta causes G0/G1 cell cycle arrest through induction of cyclin-dependent kinase inhibitors. EMBO J, 15, 2748–2759.

47. Rodriguez, A., Jung, E.J. and Flemington, E.K. (2001) Cell cycle analysis of Epstein-Barr virus-infected cells following treatment with lytic cycle-inducing agents. J Virol, 75, 4482–4489.

48. Paladino, P., Marcon, E., Greenblatt, J. and Frappier, L. (2014) Identification of herpesvirus proteins that contribute to G1/S arrest. J Virol, 88, 4480–4492.

49. Ersing, I., Nobre, L., Wang, L.W., Soday, L., Ma, Y., Paulo, J.A., Narita, Y., Ashbaugh, C.W., Jiang, C., Grayson, N.E. et al. (2017) A Temporal Proteomic Map of Epstein-Barr Virus Lytic Replication in B Cells. Cell Rep, 19, 1479–1493.

50. Chiu, Y.F., Sugden, A.U. and Sugden, B. (2013) Epstein-Barr viral productive amplification reprograms nuclear architecture, DNA replication, and histone deposition. Cell Host Microbe, 14, 607–618.

51. Resnitzky, D. and Reed, S.I. (1995) Different roles for cyclins D1 and E in regulation of the G1-to-S transition. Mol Cell Biol, 15, 3463–3469.

52. Fagundes, R. and Teixeira, L.K. (2021) Cyclin E/CDK2: DNA Replication, Replication Stress and Genomic Instability. Front Cell Dev Biol, 9, 774845.

53. Pellarin, I., Dall’Acqua, A., Favero, A., Segatto, I., Rossi, V., Crestan, N., Karimbayli, J., Belletti, B. and Baldassarre, G. (2025) Cyclin-dependent protein kinases and cell cycle regulation in biology and disease. Signal Transduct Target Ther, 10, 11.

54. Honda, R., Lowe, E.D., Dubinina, E., Skamnaki, V., Cook, A., Brown, N.R. and Johnson, L.N. (2005) The structure of cyclin E1/CDK2: implications for CDK2 activation and CDK2-independent roles. EMBO J, 24, 452–463.

55. Wang, Y., Yu, J. and Pei, Y. (2024) Identifying the key regulators orchestrating Epstein-Barr virus reactivation. Front Microbiol, 15, 1505191.

56. Indari, O., Ghosh, S., Bal, A.S., James, A., Garg, M., Mishra, A., Karmodiya, K. and Jha, H.C. (2024) Awakening the sleeping giant: Epstein-Barr virus reactivation by biological agents. Pathog Dis, 82.

57. Desimio, M.G., Covino, D.A., Cancrini, C. and Doria, M. (2024) Entry into the lytic cycle exposes EBV-infected cells to NK cell killing via upregulation of the MICB ligand for NKG2D and activation of the CD56(bright) and NKG2A(+)KIR(+)CD56(dim) subsets. Front Immunol, 15, 1467304.

58. Nguyen, T.D., Wang, J., Lam, M.T., McFerrin, H., O’Grady, T.M., Roberts, C., Van Otterloo, N., Nguyen, T.T., Baddoo, M., Wyczechowska, D., et al. (2025) Comprehensive resolution and classification of the Epstein Barr virus transcriptome. Nat Commun, 16, 6381.

59. Velazquez, L., Mogensen, K.E., Barbieri, G., Fellous, M., Uze, G. and Pellegrini, S. (1995) Distinct domains of the protein tyrosine kinase tyk2 required for binding of interferon-alpha/beta and for signal transduction. J Biol Chem, 270, 3327–3334.

60. Krishnan, K., Pine, R. and Krolewski, J.J. (1997) Kinase-deficient forms of Jak1 and Tyk2 inhibit interferon alpha signaling in a dominant manner. Eur J Biochem, 247, 298–305.

61. Yiu, S.P.T., Zerbe, C., Vanderwall, D., Huttlin, E.L., Weekes, M.P. and Gewurz, B.E. (2023) An Epstein-Barr virus protein interaction map reveals NLRP3 inflammasome evasion via MAVS UFMylation. Mol Cell, 83, 2367–2386 e2315.

62. Collart, M.A., Panasenko, O.O. and Nikolaev, S.I. (2013) The Not3/5 subunit of the Ccr4-Not complex: a central regulator of gene expression that integrates signals between the cytoplasm and the nucleus in eukaryotic cells. Cell Signal, 25, 743–751.

63. Kruk, J.A., Dutta, A., Fu, J., Gilmour, D.S. and Reese, J.C. (2011) The multifunctional Ccr4-Not complex directly promotes transcription elongation. Genes Dev, 25, 581–593.

64. Villanyi, Z., Ribaud, V., Kassem, S., Panasenko, O.O., Pahi, Z., Gupta, I., Steinmetz, L., Boros, I. and Collart, M.A. (2014) The Not5 subunit of the ccr4-not complex connects transcription and translation. PLoS Genet, 10, e1004569.

65. Collart, M.A. (2016) The Ccr4-Not complex is a key regulator of eukaryotic gene expression. Wiley Interdiscip Rev RNA, 7, 438–454.

66. Hong, G.K., Delecluse, H.J., Gruffat, H., Morrison, T.E., Feng, W.H., Sergeant, A. and Kenney, S.C. (2004) The BRRF1 early gene of Epstein-Barr virus encodes a transcription factor that enhances induction of lytic infection by BRLF1. J Virol, 78, 4983–4992.

67. Hagemeier, S.R., Barlow, E.A., Kleman, A.A. and Kenney, S.C. (2011) The Epstein-Barr virus BRRF1 protein, Na, induces lytic infection in a TRAF2- and p53-dependent manner. J Virol, 85, 4318–4329.

68. Yoshida, M., Watanabe, T., Narita, Y., Sato, Y., Goshima, F., Kimura, H. and Murata, T. (2017) The Epstein-Barr Virus BRRF1 Gene Is Dispensable for Viral Replication in HEK293 cells and Transformation. Sci Rep, 7, 6044.

69. Raisch, T. and Valkov, E. (2022) Regulation of the multisubunit CCR4-NOT deadenylase in the initiation of mRNA degradation. Curr Opin Struct Biol, 77, 102460.

70. Zhu, X., Cruz, V.E., Zhang, H., Erzberger, J.P. and Mendell, J.T. (2024) Specific tRNAs promote mRNA decay by recruiting the CCR4-NOT complex to translating ribosomes. Science, 386, eadq8587.

71. Chen, Y., Boland, A., Kuzuoglu-Ozturk, D., Bawankar, P., Loh, B., Chang, C.T., Weichenrieder, O. and Izaurralde, E. (2014) A DDX6-CNOT1 complex and W-binding pockets in CNOT9 reveal direct links between miRNA target recognition and silencing. Mol Cell, 54, 737–750.

72. Wakiyama, M. and Takimoto, K. (2022) N-terminal Ago-binding domain of GW182 contains a tryptophan-rich region that confer binding to the CCR4-NOT complex. Genes Cells, 27, 579–585.

73. Bulbrook, D., Brazier, H., Mahajan, P., Kliszczak, M., Fedorov, O., Marchese, F.P., Aubareda, A., Chalk, R., Picaud, S., Strain-Damerell, C. et al. (2018) Tryptophan-Mediated Interactions between Tristetraprolin and the CNOT9 Subunit Are Required for CCR4-NOT Deadenylase Complex Recruitment. J Mol Biol, 430, 722–736.

74. Pekovic, F., Lai, W.S., Corbo, J., Hicks, S.N., Luke, K., Blackshear, P.J. and Valkov, E. (2025) Multivalent interactions with CCR4-NOT and PABPC1 determine mRNA repression efficiency by tristetraprolin. Nat Commun, 16, 7528.

75. Inada, T. and Makino, S. (2014) Novel roles of the multi-functional CCR4-NOT complex in post-transcriptional regulation. Front Genet, 5, 135.

76. Chalabi Hagkarim, N. and Grand, R.J. (2020) The Regulatory Properties of the Ccr4-Not Complex. Cells, 9.

